# Scalable continuous evolution of genes at mutation rates above genomic error thresholds

**DOI:** 10.1101/313338

**Authors:** Arjun Ravikumar, Garri A. Arzumanyan, Muaeen K.A. Obadi, Alex A. Javanpour, Chang C. Liu

## Abstract

Directed evolution is a powerful approach for engineering biomolecules and understanding adaptation^1-3^. However, experimental strategies for directed evolution are notoriously low-throughput, limiting access to demanding functions, multiple functions in parallel, and the study of molecular evolution in replicate. Here, we report OrthoRep, a yeast orthogonal DNA polymerase-plasmid pair that stably mutates ~100,000-fold faster than the host genome *in vivo*, exceeding error thresholds of genomic replication that lead to single-generation extinction^4^. User-defined genes in OrthoRep continuously and rapidly evolve through serial passaging, a highly scalable process. Using OrthoRep, we evolved drug resistant malarial DHFRs 90 times and uncovered a more complex fitness landscape than previously realized^5-9^. We find rare fitness peaks that resist the maximum soluble concentration of the antimalarial pyrimethamine – these resistant variants support growth at pyrimethamine concentrations >40,000-fold higher than the wild-type enzyme can tolerate – and also find that epistatic interactions direct adaptive trajectories to convergent outcomes. OrthoRep enables a new paradigm of routine, high-throughput evolution of biomolecular and cellular function.

---

By subjecting genes to repeated cycles of mutation and functional selection, directed evolution has yielded extraordinary successes, including numerous industrial enzymes and therapeutic proteins, expanded genetic codes, and significant insights into how RNAs and proteins evolve^1-3^. However, existing approaches to directed evolution are difficult to scale: classical methods rely on onerous rounds of *in vitro* gene diversification followed by transformation into cells for expression and selection, and a pioneering phage-assisted continuous evolution system requires specialized setups and is largely incompatible with selections based on cellular phenotypes^10^. These shortcomings limit the routine evolution of truly novel biomolecular functions that require long mutational paths to access, the rapid evolution of enzymes and metabolic pathways that fully integrate with host systems, and the extensive parallelization of directed evolution experiments to discover multiple related functions or to map the scope of adaptive trajectories leading to important functions such as drug resistance.

In principle, the most scalable and experimentally straightforward evolution systems are living cells, since populations of cells will continuously adapt when simply passaged under selective conditions. For example, microbial evolution experiments are routinely run in high-throughput (*i.e.* scores of replicate lines) to optimize strains, map adaptive landscapes, and understand evolutionary dynamics^11-16^. However, because genomic mutation rates are low, any single gene can only evolve slowly, making the basic passaging of cells a poor approach to the directed evolution of novel biomolecules or specific genes. While it is possible to increase genomic mutation rates by engineering host DNA polymerases and mismatch repair systems^4,17-19^, the high number of essential genes in a cell’s genome sets both soft and hard “speed limits” on mutation rates^4, 20-22^. Furthermore, the lack of mutational targeting would allow adaptation to occur through changes outside user-defined genes and biomolecules.

We report OrthoRep, a highly-error-prone orthogonal DNA polymerase-DNA plasmid pair^23^ that mutates user-defined genes at rates of ~1×10^−5^ substitutions per base (s.p.b.) without increasing the genomic mutation rate (~10^−10^ s.p.b.) in yeast. This ~100,000-fold mutational acceleration crosses the mutation-induced extinction threshold of the host genome^4^ to enable the rapid continuous evolution of genes entirely *in vivo* through basic serial passaging alone. We describe a substantial DNA polymerase (DNAP) engineering effort leading to highly error-prone orthogonal DNAP variants that drive the rapid evolution of only user-defined genes. We demonstrate the utility of OrthoRep in a yeast model of *Plasmodium falciparum* DHFR^24^ by evolving resistance to the maximum soluble concentration of the antimalarial DHFR inhibitor, pyrimethamine, in 90 independent 0.5 mL cultures passaged just 13 times. Prevailing analyses of *Pf*DHFR resistance focus on a single fitness peak observed widely in the field^5-9^, but our experiment accesses other peaks of equal fitness. We find that a highly adaptive first-step mutation constrains path choice through sign epistasis, leading to convergence, but also that rare mutations direct trajectories to an alternative smooth fitness peak, illustrating the balance between fate and chance in drug resistance. By drastically scaling and simplifying directed evolution, OrthoRep should have widespread utility in the search for new biomolecular and cellular functions and the study of molecular adaptation.

## Engineering highly error-prone orthogonal DNAPs for OrthoRep

The basis of OrthoRep is a DNA polymerase (TP-DNAP1) that replicates a linear, high-copy, cytoplasmic DNA plasmid, p1, in *Saccharomyces cerevisiae* (**Figure 1A**, see **Extended Data Figure 1** for a full description of OrthoRep). Owing to the unique mechanism of p1 replication initiation via protein-priming and the spatial separation of p1 and TP-DNAP1 from nuclear DNA, engineered changes to TP-DNAP1’s properties should not affect the host genome. For example, we previously found that expression of TP-DNAP1 (Y427A) moderately increased the mutation rate of genes encoded on p1 to 3.5×10^−8^ s.p.b., while the endogenous mutation rate of genes in the genome (~10^−10^ s.p.b.) remained the same ^23^.

**Figure 1.**
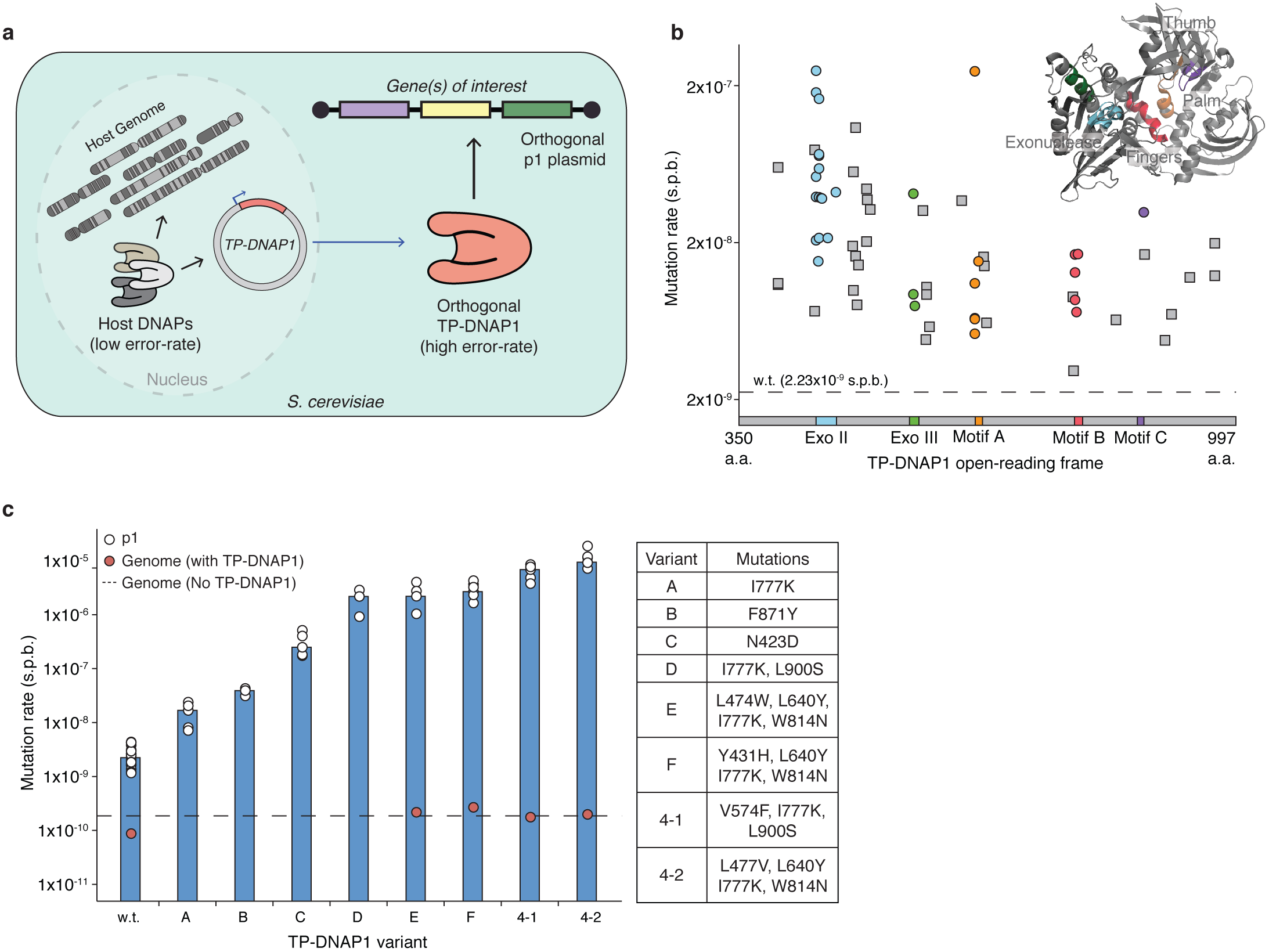
Overview and engineering of OrthoRep. **a**, Conceptual illustration of OrthoRep. **b**, Mutation rates of 65 basis set TP-DNAP1 variants found from a homology study and a TP-DNAP1 library screen. Variants are ordered by amino acid position in the TP-DNAP1 open-reading frame. Residues 1-350, corresponding to the putative TP domain, are not shown. Color-coding indicates regions known to determine fidelity. The TP-DNAP1 structure is unknown, so the closely related ϕ29 DNAP structure (PDB: 1XHX) is shown for reference with the “right hand” architecture labeled. **c**, Mutation rates of a representative panel of TP-DNAP1s and genomic substitution rates in the presence of highly error-prone variants. TP-DNAP1 substitution rates shown in **b** and **c** were measured with fluctuation tests using p1-encoded *leu2 (Q180*)*. Open circles in **c** represent measurements from independent fluctuation tests, and bars denote median measurements. Genomic substitution rates shown in **c** were determined for strains harboring p1 and each TP-DNAP1 variant as well as for the OrthoRep parent strain, AH22, which lacks p1 and TP-DNAP1. Genomic substitution rates were measured at the *URA3* locus in large-scale fluctuation tests and are shown as individual measurements. See **Extended Data Table 1** and **Supplementary Table 2** for all mutation rate values, confidence intervals, and other information.

In order to enable accelerated directed evolution experiments in small culture volumes amenable to high-throughput serial passaging, we drastically increased the mutation rate of p1 replication by engineering highly error-prone variants of TP-DNAP1. First, we found a collection of single amino acid mutant TP-DNAP1s with elevated mutation rates. We anticipated that single amino acid changes would yield modest mutators that could then serve as a basis set for building TP-DNAP1s with multiple mutations that act together to reduce fidelity. Indeed, in a study of *Escherichia coli* Pol I, a combination of three moderately fidelity-reducing mutations in the exonuclease domain, active site, and O-helix resulted in an 80,000-fold increase in Pol I’s error rate^17^. Initial attempts to populate the TP-DNAP1 basis set, based on homology analysis to related family B DNAPs, (**Extended Data Figure 2, Supplementary Table 1**) mostly yielded low activity variants (**Supplementary Table 2**), due to the idiosyncrasies of protein-priming DNAPs. Mutators identified from this effort (referred to below as Rd1 mutants) seeded our basis set, but we pursued a comprehensive approach to find high-activity variants suitable for combination.

In the largest fidelity screen of a DNAP to date, ~14,000 clones from a scanning saturation mutagenesis library of TP-DNAP1 were assayed for p1 replication activity and mutation rate, from which a set of active mutators was found. To construct the TP-DNAP1 library, we obtained an oligonucleotide pool designed to evenly sample every single-substitution amino acid variant of TP-DNAP1 without degeneracy. Oligos were designed as 29 sets, each diversifying a 20-50 amino acid region flanked by ~25 bp constant regions (**Extended Data Figure 3**). We amplified each oligo set and assembled them into full-length TP-DNAP1 variants, yielding 29 sub-libraries. Plasmid sub-libraries were screened in the OrthoRep strain, OR-Y24. OR-Y24 contains recombinant p1 that lacks wild-type *TP-DNAP1*, and instead, encodes a standardized fluorescence reporter of p1 copy number (**Supplementary Table 3**), and a disabled version of the *LEU2* selection marker (*leu2* (*Q180\**)), which serves as a reporter of substitution mutation rate in fluctuation tests that measure reversion to functional *LEU2* (see **Methods** for details). We first implemented a weak selection for TP-DNAP1 activity in OR-Y24 to eliminate frame-shifted TP-DNAP1 variants, which were common due to errors in oligonucleotide synthesis (**Extended Data Figure 4**). Purified yeast sub-libraries were plated on solid media and arrayed at 1-fold coverage, totaling 13,625 clones. (Sub-libraries 1-10, corresponding to the putative N-terminal TP of TP-DNAP1, which should not influence fidelity, were omitted.) All clones were expanded in quadruplicate and subject to p1 copy number and mutation rate measurements, using OR-Y24’s p1-encoded reporters. We carried forward 95 promising candidates and measured their mutation rates more accurately through large-scale fluctuation tests. From this, we identified 41 unique variants (Rd2 mutants) with error rates as high as ~2×10^−7^ s.p.b. (**Supplementary Table 2**). Unlike Rd1 mutants, Rd2 mutants retained high activity, and on average replicated p1 at only a 2-fold lower copy number than did wild type (w.t.) TP-DNAP1. Only 9 of the Rd2 hits contained mutations at positions considered in the homology-based library design that generated Rd1 hits, indicating that fidelity determinants of TP-DNAP1 can lie outside of the most-conserved regions of DNAPs. Incidentally, we also discovered 210 TP-DNAP1 variants that replicated p1 at a higher copy number than did w.t. TP-DNAP1 (**Supplementary Table 4**), and added the mutation from one of these variants to several low-activity mutator TP-DNAP1s to confirm the generality of the activity-boosting phenotype (**Supplementary Table 5**). These variants were not included in subsequent experiments here, but should prove useful in future TP-DNAP1 engineering efforts. Rd2 hits were combined with Rd1 hits to form a 65-member basis set (**Figure 1B**).

From the basis set mutations, we cloned and screened combinatorial libraries in two stages and found highly error-prone TP-DNAP1s. To limit combinatorial diversity, we grouped basis set mutants according to the universal DNAP “right hand” architecture^25^ and made only inter-group combinations. The structural elements responsible for DNAP fidelity primarily reside within the A and C motifs in the palm domain, B motif in the fingers domain, and Exo I, II, and III motifs in the exonuclease domain (**Figure 1B, Extended Data Figure 2**). Many of our basis set mutations lie within or near these motifs (**Figure 1B**). We expected that synergy between motif mutations from different domains (*e.g.* motif A × B) would yield super-additive or super-multiplicative reductions in fidelity, as observed with RB69 DNAP and *E. coli* Pol I, respectively^17,26^. Mutants were pooled based on proximity to motifs (see **Methods**) and distinct pools were shuffled with each other in two rounds of screening. We found 46 mutators by screening a library of motif B mutants crossed with motif A and C mutants, including three TP-DNAP1s (Rd3 mutants) with mutation rates of ~1×10^−6^ s.p.b. (9.70×10^−7^-3.36×10^−6^ s.p.b., 1.35×10^−6^ s.p.b., 9.22×10^−7^-2.87×10^−6^ s.p.b., see **Supplementary Table 2**), representing a ~400-fold increase over the w.t. TP-DNAP1 mutation rate and a ~10,000-fold increase over the genomic mutation rate. Mutations at I777 in motif B were broadly synergistic in combination with mutations in motifs A and C and produced super-multiplicative increases in error rate (**Supplementary Table 2**). Rd3 mutants were crossed with all of the exonuclease mutants from the basis set in a final round of screening. This yielded two highly error-prone variants, TP-DNAP-4-1 (V574F, I777K, L900S) and TP-DNAP1-4-2 (L477V, L640Y, I777K, W814N), with mutation rates of ~7×10^−6^ s.p.b. (3.78×10^−6^-8.53×10^−6^ s.p.b.) and ~1×10^−5^ s.p.b. (7.19×10^−6^ -1.88×10^−5^ s.p.b.), respectively (**Figure 1C, Supplementary Table 2**). For facile generation of DNA libraries *in vivo* with TP-DNAP1-4-2, a 0.3 μL saturated yeast culture is theoretically sufficient to achieve 1-fold coverage of all single mutants of a 1 kb gene and a 90 mL culture is sufficient for all double mutants. Moreover, high p1 mutation rates remained completely stable for the longest duration tested (90 generations; **Supplementary Table 2**) and genomic mutation rates remain unchanged in the presence of p1 replication by TP-DNAP1-4-2 (**Figure 1C, Extended Data Table 1**), meaning that OrthoRep can sustain *in vivo* mutagenesis with complete orthogonality (*i.e.* at least ~100,000-fold mutational targeting) for continuous evolution experiments.

## Crossing the mutation-induced extinction threshold of the yeast genome

OrthoRep can access and sustain mutation rates that untargeted genome mutagenesis cannot. There is a theoretically predicted and empirically observed inverse relationship between the length of an information-encoding polymer, such as a gene or genome, and the tolerable error rate of replication^4,20,21^. At sufficiently high mutation rates, essential genetic information is destroyed every generation, guaranteeing extinction; and even moderately elevated mutation rates can cross intermediate “error thresholds” where replicative fitness erodes^22^. OrthoRep can bypass the low extinction and error thresholds of large cellular genomes *in vivo* by targeting mutations to just p1, which contains only genes of interest. In contrast, continuous evolution systems that use whole-genome mutagenesis of cells or phages are still subject to the lower thresholds of the host or phage genomes, respectively^10, 17-19^. We compared targeted mutagenesis via OrthoRep to whole-genome mutagenesis by experimentally applying high mutation rates to the host genome. This was done by transplanting a subset of mutations, discovered by Herr *et al.*^4^, that increase the substitution mutation rate of POL3, the primary yeast lagging strand DNAP, into wild-type or mismatch repair-deficient (*Δmsh6*) versions of AH22, the parent of OR-Y24. Mutator phenotypes (verified by fluctuation tests at a genomic locus) were accompanied by severe growth defects and lead to single-generation extinction in the case of *pol3-01*, *Δmsh6* AH22 (**Figure 2A**). In accordance with Herr *et al.*^4^, the projected mutation rate imposed in this nonviable strain is 4.72×10^−6^ s.p.b., calculated from the individual contribution of *pol3-01* and the average effect of *MSH6* loss. We interpret this mutation rate to be above the extinction threshold, causing an error catastrophe that kills the host cell. Since replication of p1 by TP-DNAP1-4-2 occurs at a higher mutation rate than 4.72×10^−6^ s.p.b., we conclude that OrthoRep can stably exceed categorical mutation rate limits on replicating cellular genomes. We also asked whether viable genomic mutator strains could sustain mutagenesis. Four AH22 strains with mutation rates of 1.64×10^−7^ -5.24×10^−7^ s.p.b. were propagated for 82 generations in triplicate and afterwards, a clone from each was subject to genomic mutation rate measurements via fluctuation tests (**Figure 2B**). Across replicates, the mutation rate drops an average of 284-fold, indicating that in durations relevant to directed evolution experiments, continuous genome mutagenesis is unsustainable whereas continuous mutagenesis of p1 is sustainable (**Figure 2B**).

**Figure 2.**
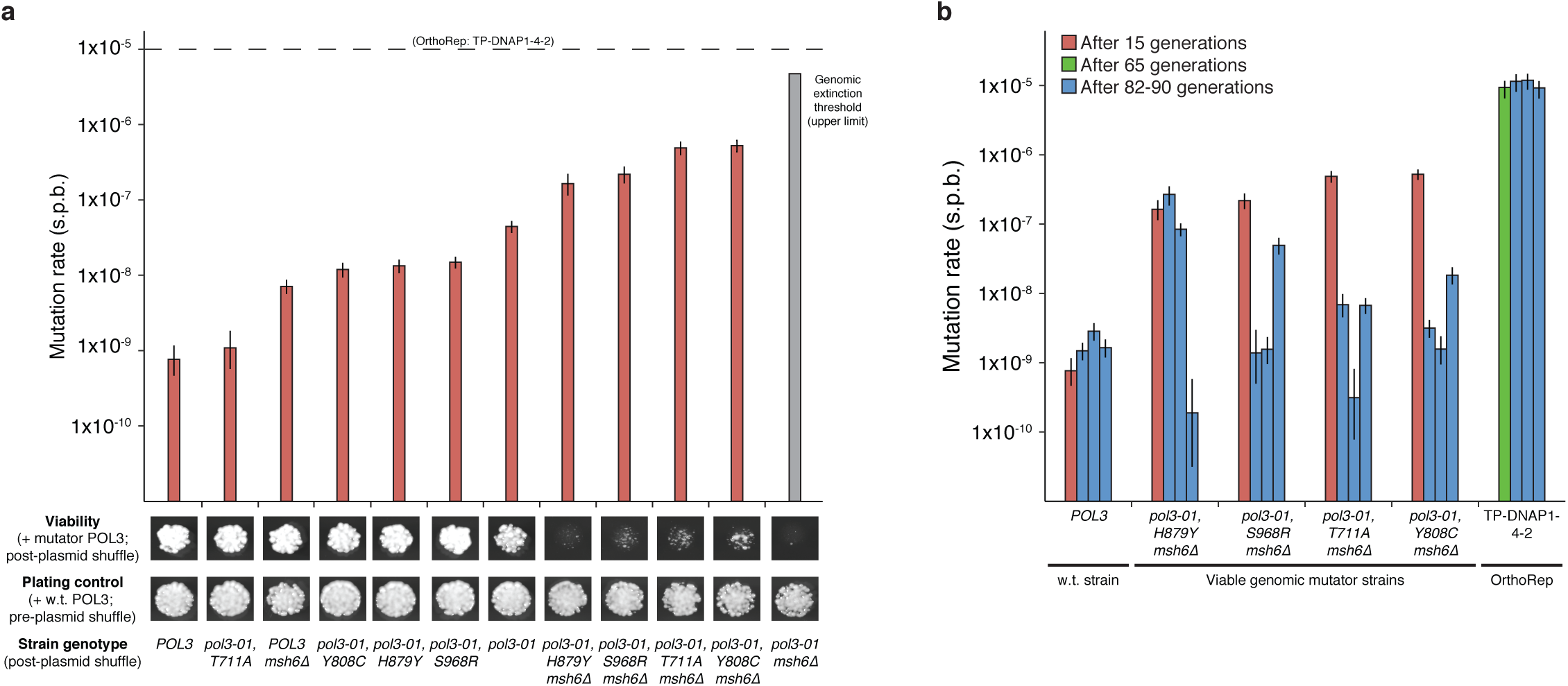
Mutation-induced extinction and error thresholds of the yeast genome. **a**, A series of yeast genomic mutator strains spanning mutation rates from the w.t. rate to the extinction threshold (upper limit). OrthoRep’s parent strain, AH22, was modified to express its genomic w.t. *POL3* from a plasmid, and *POL3* variants were introduced into w.t. or mismatch repair-deficient (Δ*msh6*) versions of this strain *via* plasmid shuffle. W.t. *POL3* is retained in pre-plasmid shuffle plating controls. Genomic mutation rates were measured ~15 generations after plasmid shuffle. The projected mutation rate of the inviable *pol3-01,* Δ*msh6* strain was calculated as the product of the mutational increases due to pol3-01 (58-fold) and Δ*msh6* mutations (106-fold, averaged across genotypes). The proofreading deficient *pol3-01* allele encodes *POL3* (*D321A, E323A*). T711A, Y808C, H879Y, and S968R are suppressor mutations that reduce the error-rate of *pol3-01*. **b**, Mutational stability of viable genomic mutator strains versus OrthoRep. Strains harboring *POL3* variants or OrthoRep with TP-DNAP1-4-2 were passaged in triplicate for 82 or 90 generations, respectively. Afterwards, genomic or OrthoRep substitution mutation rates were measured at the genomic *CAN1* locus or with p1-encoded *leu2 (Q180*)*, respectively. Data shown in **a** and **b** are individual measurements with 95% confidence intervals.

## High-throughput evolution of PfDHFR to understand adaptive trajectories leading to drug resistance

Sustainable, continuous, and targeted mutagenesis with OrthoRep can be used to understand and predict drug resistance in high-throughput evolution experiments that abundantly sample adaptive trajectories and outcomes. *Pf*DHFR resistance to the antimalarial drug, pyrimethamine, occurs in the wild primarily through four active site mutations (N51I, C59R, S108N, and I164L), but the broader resistance landscape remains largely unknown. Laboratory evolution and landscape-mapping studies have mostly been limited to the quadruple mutant fitness peak (qm-wild) and suggest that resistance reproducibly arises from the crucial S108N mutation, followed by step-wise paths to qm-wild^5-9, 24^. We asked whether high-throughput directed evolution of *Pf*DHFR resistance to pyrimethamine would reveal a more complex landscape with additional fitness peaks, including ones that forgo S108N.

We applied OrthoRep in a well-established yeast model of *Pf*DHFR^24^ and evolved resistance to pyrimethamine in 90 independent 0.5 mL cultures (**Figure 3A**). Transgenic yeast strains that lack endogenous DHFR and depend on p1-encoded *Pf*DHFR acquired sensitivity to pyrimethamine and in pilot studies, were able to evolve resistance by mutating *Pf*DHFR (**Extended Data Figure 5**). OR-Y8, which uses TP-DNAP1-4-2 to replicate p1 encoding w.t. *Pf*DHFR, was used to seed 90 independent 0.5 mL cultures containing pyrimethamine. Cultures were grown to saturation and uniformly passaged at 1:100 dilutions into media containing gradually increasing pyrimethamine concentrations chosen to maintain strong selection as populations adapted (**Figure 3A**). After just 13 passages (*i.e.* 87 generations), 78 surviving populations adapted to media containing the maximum soluble concentration of pyrimethamine (3 mM). (Extinction was stochastic in duplicate revival experiments (**Supplementary Table 6**).) From Sanger sequencing analysis (see **Methods** for details) of bulk adapted populations, we identified 37 unique *PfDHFR* coding mutations across all replicates and as many as six amino acid changes in a single population (replicates 16 and 61 in **Figure 3B**). Ten of these 37 mutations have been previously reported to yield pyrimethamine resistance^7-9, 27^. Several mutations identified in the promoter region increased gene expression (unpublished results) and we hypothesize that some of the observed synonymous mutations in *PfDHFR* reduce translational suppression mediated by binding of *Pf*DHFR to its own mRNA sequence^28^.

**Figure 3.**
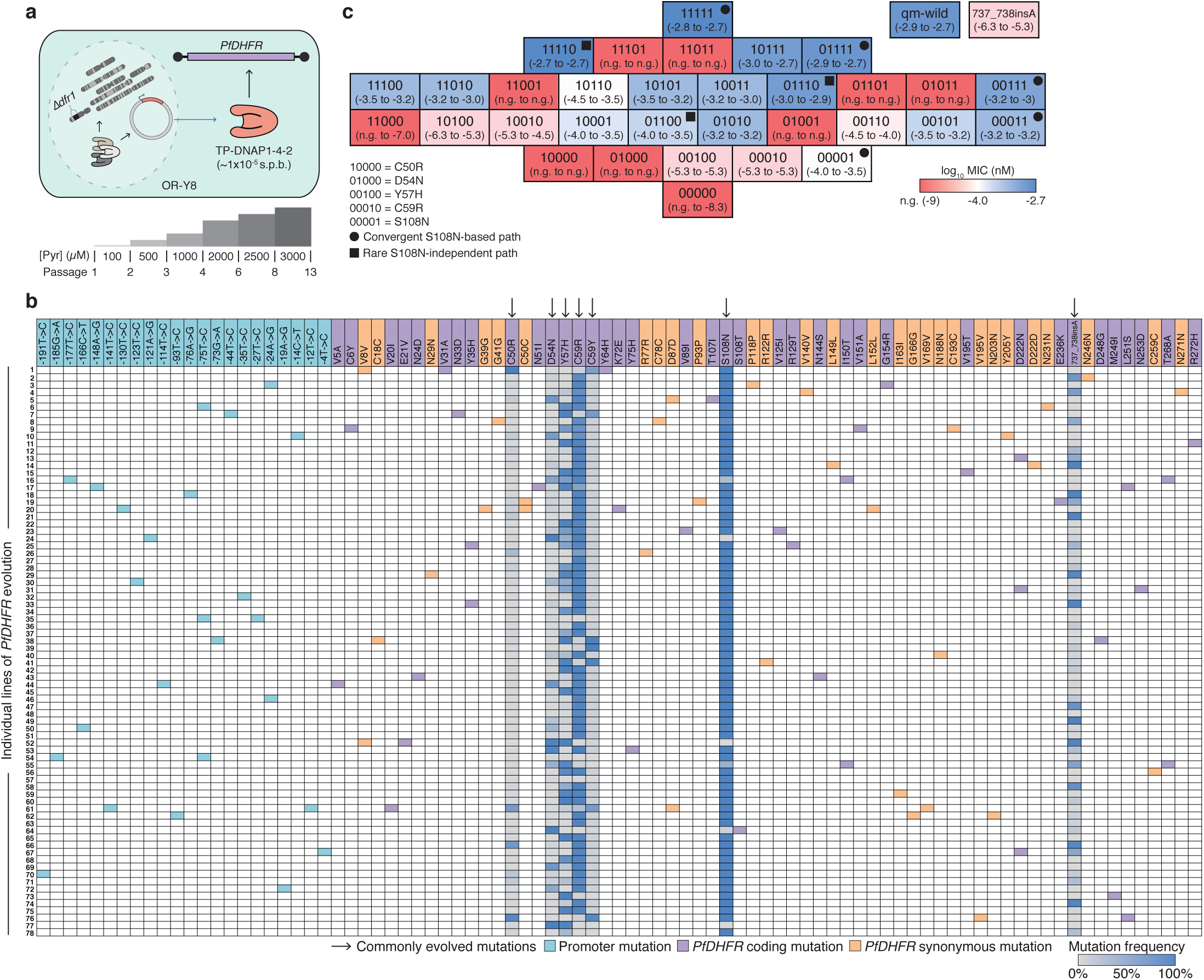
High-throughput evolution of PfDHFR resistance to pyrimethamine. **a**, The strain used for evolution (OR-Y8) and the drug regimen to which it was subjected. Evolving lines of OR-Y8 were monitored daily by OD_600_ measurement and passaged at a 1:100 dilution when 80/90 replicates reached an OD_600_ of 0.7. Pyrimethamine concentration was uniformly increased if diluted cultures reached the growth cutoff within 48 hours. Evolution was terminated when populations fully adapted to 3 mM pyrimethamine. **b**, *PfDHFR* and promoter mutations identified in 78 evolved populations from Sanger sequencing. Green, purple, and yellow shading indicates the presence of a mutation at ~20% frequency or higher. For seven commonly observed mutations, frequencies were calculated and are shown with a color on a gray-blue scale. See **Methods** for SNP analysis details. **c**, A fitness map of a five-mutation *PfDHFR* landscape defined by C50R, D54N, Y57H, C59R, and S108N. MIC of pyrimethamine was determined for yeast strains expressing all 32 *PfDHFR* alleles from this landscape. Data shown are the range of log_10_(MIC of pyrimethamine (nM)) for biological triplicates, with a color on a red-blue scale indicating the median. The mid-point of the red-blue scale is shifted to distinguish highly resistant alleles. n.g., no growth.

Adapted populations primarily converged on a previously unidentified S108N-based fitness peak equal in fitness to qm-wild. Across all replicates, we observed seven pervasive coding changes (**Figure 3B**), including 737_738insA, which creates an adaptive truncation of the C-terminal linker region (**Figure 3C**). Sixty-two of the 78 adapted populations had high frequencies of two mutations present in qm-wild (C59R and S108N), but only one of these populations accumulated a third mutation (N51I) from the qm-wild peak (replicate 17 in **Figure 3B**). Instead, most populations carried combinations of C50R, D54N or Y57H in addition to C59R and S108N, and two populations contained D54N, Y57H, C59R, and S108N together (replicates 16 and 30 in **Figure 3B**), indicative of a new fitness peak. To validate this, we mapped resistance of a five-mutation landscape defined by C50R, D54N, Y57H, C59R, and S108N (**Figure 3C**). (For characterization, *PfDHFR*s were cloned into a nuclear plasmid to avoid rapid mutation and expressed at low levels to amplify differences between highly resistant alleles.) We found that the quadruple mutant peak reached in our experiment (01111, see **Figure 3C**) has the same pyrimethamine MIC as qm-wild, demonstrating that OrthoRep can discover new, highly adaptive alleles. Moreover, 11111 is more resistant than qm-wild, and would have likely been observed in our replicate populations if stronger pyrimethamine selection were feasible.

Epistasis among mutations in S108N-based trajectories directs adaptation to 11111 and leads to the observed convergence of 00111 and 01111 intermediates across replicate lines. Because S108N is a highly adaptive single mutant, 00001 rapidly and repeatedly fixed first in evolving populations (**Extended Data Figure 6**), and blocked access to the 96/120 possible trajectories to 11111 that start with other first-step mutations. From 00001, access to 11111 is additionally constrained by negative epistasis between S108N and D54N, which is relieved and changes sign only when Y57H and C59R are both present (**Figure 3C**). (We note that adapted populations in our evolution experiment containing high frequencies of D54N, C59R and S108N without Y57H, typically carry other, potentially compensatory, promoter and coding mutations (**Figure 3B**)). As a result, just eight of the 24 possible paths from 00001 to 11111 avoid inactive *Pf*DHFR intermediates. Of these remaining paths, the greediest one (00000→00001→00011→00111→01111→11111) likely produced the repeated evolution of populations containing 00111 and 01111.

Notably, an additional fitness peak (11110) avoids S108N. Three adapted populations lack mutation at S108 (**Figure 3B**) and can access this most-resistant quadruple mutant. We attribute this to a rare clonal interference event where the 01100 double mutant arises and displaces a population that has nearly fixed 00001 (**Extended Data Figure 6A, F**). One of these replicates additionally fixed C59R to reach the triple mutant (01110) with the highest MIC (**Figure 3B**); and additional selection pressure should fix C50R to reach 11110 (**Figure 3C**). Since 11110 is a smooth peak, weaker early selection or greater population structure^14,29^, both of which should allow adaptive first-step mutations other than S108N (*e.g.* Y57H, C59R) to fix, would increase the chance of reaching 11110 and decrease repeatability of path choice. This has implications for drug schedule design as it can dictate both the identity of resistant genotypes reached and the reproducibility of adaptive trajectories.

Several adaptive populations also access the broader landscape beyond 11111. We find a rugged local maximum containing C59Y (10121) that traps a few evolving populations (**Figure 3B, Extended Data Figure 7**). We also observe a single replicate that fixes D54N with S108T and avoids negative epistasis with S108N (**Figure 3B**). Future analysis will include less frequent putative adaptive mutations that occur in multiple replicates (*e.g.* Y35H, I150T, D222N, L251S, T268A from **Figure 3B**) or fix independently in time (*e.g.* M249I from **Extended Data Figure 6H**). However, our analysis of only the most common adaptive mutations and mutational paths has already uncovered new peaks in the landscape of *Pf*DHFR-mediated drug resistance and provides examples of how epistasis results in evolutionary repeatability and how the existence of greedy mutations such as S108N can render a highly adaptive outcome (11110) rare through early fixation. In other words, high-throughput directed evolution with OrthoRep enables the discovery of new fitness peaks and thorough studies of molecular adaptation at the level of a single protein.

## Discussion

OrthoRep should have broad utility for directed evolution due to its unique features. First, OrthoRep realizes continuous mutagenesis entirely *in vivo*. Therefore, it readily integrates with the existing rich ecosystem of yeast genetic selections including growth-based positive selections; dominant negative selections that may require titration of p1’s copy number (**Extended Data Figure 8**); selections utilizing cell-based technologies such as fluorescence-activated cell sorting (FACS), continuous culturing devices, and droplet screening systems; and selections for cell-level phenotypes such as drug resistance, stress tolerance, or metabolism^1-3, 30^. To enable its widespread application, we have also established OrthoRep in different yeast backgrounds, including diploids and industrially relevant strains (manuscript in preparation). Second, OrthoRep is a scalable directed evolution platform, since it does not require *in vitro* library construction or specialized equipment. Therefore, it can be used to evolve genes at bioreactor-scale or, as demonstrated here, in small culture volumes in a high-throughput manner with basic serial passaging. In addition to drug resistance and fitness landscape studies, large high-throughput replication of evolution experiments can be used to test the relationship between adaptive outcomes and mutational supply, gene dosage, population size, population structure, or selection dynamics. Genes can also be evolved for many related phenotypes (*e.g.* biosensors that recognize different substrates) in parallel. Third, OrthoRep supports custom mutation rates *in vivo*. Ongoing engineering of TP-DNAP1, informed by *in vitro* characterization and structure determination, should yield variants that approach the extinction threshold of a typical 1 kb gene (~10^−3^), thereby maximizing the mutation rate for continuous *in vivo* directed evolution. TP-DNAP1s can also be engineered with custom mutational spectra (**Extended Data Table 2**) or with high in/del rates for optimizing loop regions of synthetic protein scaffolds. Finally, OrthoRep is a simple and highly stable continuous evolution system that allows for the routine traversal of long mutational trajectories leading to demanding biomolecular and cellular functions.

## Acknowledgements

We thank members of our group, especially T. Loveless and Z. Zhong for helpful discussions and suggestions. We thank S. Ahrar, H. Rishi, and M. Shapiro for valuable comments on the manuscript. This research was funded by the Defense Advanced Research Projects Agency (HR0011-15-2-0031), the National Institutes of Health (1DP2GM119163-01), the National Science Foundation (MCB1545158), the Arnold and Mabel Beckman Foundation, and startup funds from UC Irvine.

## Author contributions

A.R. designed and performed TP-DNAP1 screens, error threshold experiments, and *Pf*DHFR evolution experiments. G.A.A. designed and performed *Pf*DHFR evolution experiments. M.K.A.O. assisted in TP-DNAP1 screens. A.A.J. generated homology analysis for TP-DNAP1s. A.R., G.A.A., and C.C.L. analyzed results and wrote the manuscript. C.C.L. supervised the research.

## Competing interests

C.C.L. and A.R. have filed a provisional application with the US Patent and Trademark Office on this work.

## Materials & Correspondence

All requests for materials and correspondence should be addressed to C.C.L.

## Methods

### DNA cloning

Plasmids used in this study are listed in **Supplementary Table 7**. *E. coli* strain TG1 (Lucigen) was used for all of the DNA cloning steps. All primers used in this study were purchased from IDT. All enzymes for PCR and cloning were obtained from NEB. All individually cloned plasmids (*i.e.* excluding TP-DNAP1 libraries) were assembled by the Gibson method^31^. To clone plasmid 29, a DNA fragment encoding the open-reading frame of *PfDHFR* (819 bp) was obtained from IDT.

To clone the scanning saturation mutagenesis library of TP-DNAP1, a pool of ~19,000 oligonucleotides (130-200-mers) was obtained from Agilent Technologies and sub-cloned into plasmid 2. The oligo pool was designed as 29 sub-libraries, each covering a 25-50 variable amino acid region of the TP-DNAP1 open-reading frame and flanked by ~25 bp constant regions. The variable region consisted of a replacement of each amino acid in the wild-type sequence with 19 codons representing the 19 other amino acids. The mutagenic codons were chosen from a 20-codon genetic code with a maximal codon adaptation index for the *S. cerevisiae* genome. Constant regions were chosen for efficient PCR amplification. Each sub-library was PCR amplified and assembled into corresponding PCR-amplified plasmid 2 backbones by the Gibson method^31^. Assembled sub-libraries were transformed into *E. coli* at >30-fold coverage of theoretical diversity and plated on selective LB plates. After overnight growth at 37 °C, transformants were scraped from plates and resuspended in 0.9% NaCl for plasmid extraction. Control transformations containing only the plasmid 2 backbones were similarly treated, to verify a low frequency (<5%) of full-length plasmid 2 carry-over. Plasmids were extracted from individual clones of two sub-libraries and subject to analysis via agarose gel electrophoresis and Sanger sequencing.

To clone mutant TP-DNAP1 shuffling libraries, plasmids of the 65 basis set mutants were pooled and crossed by the Gibson method^31^. Since many basis set mutations encode mutations outside of strictly conserved motifs, the TP-DNAP1 open-reading frame was segmented into four regions to define broader boundaries for shuffling: the exonuclease domain (amino acids 1-596), motif A (amino acids 597-684), motif B (amino acids 685-819) and motif C (amino acids 820-987). To cross the 7 motif B basis set mutants with the 10 motif A and 8 motif C basis set mutants, the corresponding regions were PCR amplified from individual mutant TP-DNAP1 plasmids, and PCR amplicons from each region were pooled in equimolar ratios. Pooled fragments were assembled with a PCR-amplified plasmid 2 backbone by the Gibson method^31^. Assembled libraries were transformed and extracted as described above. Shuffling libraries contained a large fraction of misassembled plasmids, as determined by agarose gel electrophoresis. The desired plasmid population was purified by gel extraction and re-transformation. Both transformation steps retained >100-fold coverage of theoretical library size. Plasmids were extracted from individual clones of the purified libraries and subject to analysis via gel electrophoresis and Sanger sequencing. To cross round-3 mutants with exonuclease basis set mutants, a new region was defined to cover round-3 mutants (amino acids 597-987), and a similar cloning procedure was followed.

### Yeast transformation and DNA extraction

All yeast transformations (including p1 integrations) and whole-cell yeast DNA extractions were performed as described previously^23^. Genomic modifications were made using a CRISPR-Cas9 system for *S. cerevisiae*^32^.

We note the following protocol modifications for library transformations: (i) 10 μg of plasmid DNA was added for each library transformation; (ii) cells were incubated at 30 °C for 45 min with rotation at ~10 r.p.m prior to heat shock; (iii) cells were resuspended in 0.9% NaCl after heat shock and a small portion was plated on selective synthetic complete (SC) medium to determine library size; (iv) the remaining resuspension was inoculated directly into 50 mL (per transformation) of selective SC media and grown to saturation at 30 °C.

### Yeast strains

Yeast parent strains used in this study are listed in **Supplementary Table 8**. Parent strains do not include OR-Y24-derived strains containing TP-DNAP1 variants or AR-Y432-derived strains containing POL3 variants. Strains AH22 and F102-2 are described previously^23^.

Strains AR-Y285, AR-Y293, AR-Y302, AR-Y401, AR-Y443, and GA-Y109 contain a mixture of w.t. p1 and recombinant p1 containing a yeast selection marker in place of *TP-DNAP1*. These strains were not propagated beyond the original outgrowth step from the transformation plate, to retain a ~50:50 mixture of w.t. and recombinant p1. Glycerol stocks of meta-stable strains stored at -80 °C served as the source for all subsequent yeast transformations of TP-DNAP1 variants.

### TP-DNAP1 homology analysis

A list of 99 homologs to TP-DNAP1 (EMBL accession number: CAA25568.1) was generated via protein BLAST^34^ with default settings (**Supplementary Table 1**). A multiple sequence alignment of TP-DNAP1 to these homologs was performed using Clustal Omega^35^ and the resulting alignment was analyzed using Jalview^36^.

Amino acid mutations were selected based on three criteria. First, candidate positions should be flanked on both sides by residues with sequence alignment to >75% of homologs. Second, the TP-DNAP1 amino acid at a candidate position should be represented across >25% of homologs. Third, amino acids not present in TP-DNAP1 at a candidate position should be conserved across >25% of homologs. If these criteria were met, then amino acids identified from the third criterion were introduced at the candidate position in TP-DNAP1.

### Functional purification of TP-DNAP1 sub-libraries

The pilot study shown in **Extended Data Figure 4** confirmed that sub-libraries transformed into OR-Y24 are enriched for full-length TP-DNAP1 variants after two passages in SC media lacking uracil and histidine (SC-UH). To purify the entire scanning saturation mutagenesis library, the remaining 27 TP-DNAP1 plasmid sub-libraries were individually transformed into OR-Y24, and the resulting yeast sub-libraries were passaged twice at 1:100 dilutions in SC-UH. Passaged yeast sub-libraries were individually plated on solid SC medium lacking histidine. For each sub-library, 24 colonies were propagated in small cultures of SC-UH, in order to verify that >90% of clones robustly grow under selection for p1 replication. Afterwards, colonies from each yeast sub-library were individually inoculated into small cultures of SC-UH at ~1-fold coverage of theoretical sub-library diversity. Arrayed yeast sub-libraries were grown to saturation, passaged in SC-UH to fully displace w.t. p1, and then subject to small-scale p1 fluctuation tests, as described below.

### Small-scale p1 fluctuation tests

Small-scale fluctuation tests of p1-encoded *leu2 (Q180*)* were performed for TP-DNAP1 library screens. As described previously^23^, *leu2 (Q180*)* contains a C→T mutation at base 538 in *LEU2* at a site permissive to all single point mutants that generate missense mutations. To screen TP-DNAP1 libraries, OR-Y24 strains were transformed with TP-DNAP1 plasmid libraries and the resulting yeast libraries were passaged in SC-UH to displace w.t. p1. Cured strains were diluted 1:10,000 into selective SC media for fluctuation tests. Selective SC media used for p1 fluctuation tests lacked uracil, histidine and tryptophan, and was adjusted to pH 5.8 with NaOH (SC-UHW, pH 5.8). Absence of tryptophan and pH adjustment inhibited growth on reversion medium resulting from nonsense suppression of *leu2 (Q180*)*. Dilutions were split into three 100 μL cultures and one 200 μL culture in 96-well trays, and cultures were grown to saturation for 2-2.5 days. Saturated 200 μL cultures were subject to a copy number measurement, as described below. The remaining three replicates were washed and resuspended in 35 μL 0.9% NaCl. 10 μL was spot plated onto solid SC medium selective for *LEU2* revertants. Solid SC medium used for p1 fluctuation tests lacked uracil, histidine, tryptophan and leucine and was adjusted to pH 5.8 with NaOH (SC-UHLW, pH 5.8). Plates were incubated at 30 °C for 5-6 days, and afterwards, colony-count was determined for each spot.

Small-scale fluctuation data were analyzed by the MSS maximum-likelihood estimator method^37^ implemented using a custom MATLAB script. Measuring cell titers was infeasible due to the large number of strains, so the average number of cells per culture was assumed to remain constant. Relative phenotypic mutation rates were calculated by normalization to p1 copy number.

In the screen of the TP-DNAP1 scanning saturation mutagenesis library, 13,625 yeast clones were subject to small-scale p1 fluctuation tests. 376 clones with the highest relative phenotypic mutation rates were subject to an additional small-scale p1 fluctuation test with six replicates. TP-DNAP1 expression vectors were isolated from 95 yeast clones with the highest relative phenotypic mutation rates and subject to Sanger sequencing. These TP-DNAP1s were characterized with large-scale p1 fluctuation tests, as described below.

Shuffling libraries were screened in a similar manner. From the first stage, 1520 yeast clones were subject to small-scale p1 fluctuation tests. 188 clones were subject to additional small-scale p1 fluctuation tests and isolated for Sanger sequencing. 46 unique variants with the highest relative phenotypic mutation rates were characterized with large-scale p1 fluctuation tests, as described below. In the second stage, 744 yeast clones were screened and 58 clones were characterized with additional small-scale p1 fluctuation tests. Four clones were extracted, subject to Sanger sequencing and characterized with large-scale p1 fluctuation tests, as described below.

### Large-scale p1 fluctuation tests

Large-scale fluctuation tests of p1-encoded *leu2 (Q180*)* were performed to measure per-base substitution rates for individually cloned or isolated TP-DNAP1 variants. Large-scale p1 fluctuation tests are performed similarly to small-scale p1 fluctuation tests, with several modifications. First, large-scale p1 fluctuation tests were typically performed with 36-48 replicates. For highly error-prone TP-DNAP1s obtained from later rounds of screening, fewer replicates (3-16) were used to achieve similar precision^37^. Second, p1 copy number was determined by the flow cytometry method, described below. Third, cell titers were measured for each fluctuation test to estimate the average number of cells per culture. Cell resuspensions were diluted and plated on solid SC-UH medium, and colony counts were determined after incubation at 30 °C for 2-3 days. Alternatively, cell resuspensions were diluted and subject to an event-count measurement via flow cytometry. Fourth, inoculums of highly error-prone TP-DNAP1s occasionally contained preexisting mutants, despite the 1:10,000 dilution, so mutant frequencies were estimated by plating precultures on solid SC-UHLW, pH 5.8 medium. Plates were incubated for 2-3 days, in parallel with cultures grown for fluctuation tests. Preculture mutant titers were counted to estimate the number of replicates in the fluctuation test expected to contain preexisting mutants (n). Revertants were counted from all replicates of the fluctuation tests, counts were sorted, and n replicates with the highest counts were omitted from calculations.

To calculate per-base substitution rates, fluctuation data were analyzed by the maximum likelihood method, implemented using newton.LD.plating in rSalvador 1.7^38^. Phenotypic mutation rates were calculated by normalizing to the average number of cells per culture. Phenotypic mutation rates were divided by the measured p1 copy number and by the number of ways *leu2 (Q180*)* can revert to *LEU2* (2.33 for the ochre codon) to yield per-base substitution rates. 95% confidence intervals were similarly scaled by these factors.

### p1 copy number determination

A calibration curve was established to correlate p1 copy number, determined via quantitative PCR (qPCR), with fluorescence of p1-encoded *mKate2*. Five TP-DNAP1 variants representing a wide range of copy numbers were transformed into OR-Y24 and passaged until w.t. p1 was displaced. The five OR-Y24 strains were grown to saturation, diluted 1:10,000 to mimic p1 fluctuation tests, and grown in triplicate 100 μL cultures and duplicate 40 mL cultures. 100 μL saturated cultures were subject to fluorescence measurement of mKate2 (ex/em = 561 nm/620 nm, bandwidth = 15 nm) on a flow cytometer (Invitrogen Attune NxT). Whole-cell DNA extracts were prepared from 40 mL cultures and used as templates for qPCR measurement of p1-encoded *leu2 (Q180*)*, as described previously^23^. The correlation of mKate2 fluorescence and qPCR-determined p1 copy number had a strong linear fit across p1 copy numbers ranging from 9-90, and had low background (y = 0.048x + 0.206, r^2^ = 0.954).

To determine p1 copy number for large-scale p1 fluctuation tests, additional replicates of OR-Y24 strains were grown from the 1:10,000 dilution in triplicate and saturated cultures were subject to fluorescence measurement via flow cytometry. Fluorescence measurements were converted to copy number with the linear calibration curve.

To determine relative p1 copy number for small-scale p1 fluctuation tests, an additional 200 μL culture of OR-Y24 strains were grown from the 1:10,000 dilution. 200 μL saturated cultures were subject to an OD_600_ measurement and fluorescence measurement of mKate2 (ex/em = 565 nm/630 nm), using a microplate reader (TECAN Infinite M200 PRO). A linear relationship was assumed between p1 copy number and OD_600_-normalized mKate2 fluorescence. Copy numbers were calculated from normalization to a w.t. TP-DNAP1 control.

For a few large-scale p1 fluctuation tests, p1 copy number was determined by the method of small-scale p1 fluctuation tests (i.e. microplate reader measurement). For these experiments, mutant TP-DNAP1 p1 copy numbers were calculated by normalization to a w.t. TP-DNAP1 control. The copy number of this control was assumed to be the average w.t. TP-DNAP1 copy number from all large-scale p1 fluctuation tests that used flow cytometry measurements.

### Characterization of high activity TP-DNAP1s

From mKate2 measurements of the 13,625 clones screened with small-scale p1 fluctuation tests, 283 variants with high p1 copy number were chosen for additional characterization. TP-DNAP1 plasmids were isolated from these strains and subject to Sanger sequencing. 210 unique variants were identified and corresponding plasmids were re-transformed into OR-Y24. Transformed strains were passaged in SC-UH until w.t. p1 was displaced, and subject to p1 copy number measurement in triplicate. Four high copy variants were additionally confirmed by qPCR measurements (unpublished results). To test the suppressor activity of the G410H mutation, which yields increased p1 copy number, this amino acid change was added to several low-activity TP-DNAP1 variants. p1 copy number was determined for these strains, as described previously.

### Characterization of mutational preferences of TP-DNAP1s *in vivo*

Two additional fluctuation tests, coupled with sequencing, were used to determine mutational preferences of TP-DNAP1s, as described previously^23^. In one experiment, strain AR-Y302 was used for fluctuation tests. AR-Y302 contains recombinant p1 encoding *leu2 (Q180*)*, which contains two substitutions (538C>T and 540A>G), which create an amber nonsense mutation (TAG) in *LEU2*. Fluctuation tests can be used to determine the frequency of reversion from *leu2 (Q180*)* to *LEU2*. Single point mutations that restore *LEU2* from *leu2 (Q180*)* include T:A→A:T, T:A→C:G, T:A→G:C, G:C→T:A and G:C→C:G. This leaves the G:C→A:T mutation unrepresented. To measure the rate of this final mutation, we used strain AR-Y401, which contains recombinant p1 encoding *ura3 (K93R)*. This disabled allele contains a 278A>G substitution and the only way to restore *URA3* is via the G:C→A:T mutation^23^.

Strains of AR-Y302 containing TP-DNAP1 variants were subject to large-scale p1 fluctuation tests. p1 plasmids were extracted from 50-70 revertant colonies, each from an independent replicate of the fluctuation test. The restored *LEU2* alleles were PCR amplified and subject to Sanger sequencing. To calculate individual per-base pair mutation rates, the corresponding per-base substitution rate (measured from fluctuation tests of the corresponding TP-DNAP1 in OR-Y24) was used. Per-base substitution rates calculated from reversion of *leu2 (Q180*)* is more specifically the sum of the mutation rates of T:A→A:T, T:A→C:G, and T:A→G:C, weighted in proportion to the number of possible sites at which each mutation can occur and be detected. The relative preferences for these mutations, determined from Sanger sequencing, was used to calculate per-base pair mutation rates.

The G:C→A:T mutation rate is equal to the per-base mutation rate calculated from fluctuation tests of TP-DNAP1s expressed in AR-Y401. Fluctuation tests were performed and analyzed by the large-scale protocol described above. The G:C→A:T mutation rate was proportionally added to the full substitution spectra.

### Genomic fluctuation tests in the presence of error-prone TP-DNAP1s

Fluctuation tests of the genomic *URA3* gene were performed to determine genomic per-base substitution rates, as previously described^23^. Fluctuation data were analyzed by the maximum likelihood method, as described for large-scale p1 fluctuation tests. Phenotypic mutation rates were divided by the target size for loss of function of *URA3* via base pair substitution^39^, to yield per-base substitution rates. 95% confidence intervals were similarly scaled by these factors.

### Construction and characterization of *POL3* mutator strains

To construct *POL3* mutator strains, plasmids 19-25 were transformed into AR-Y432 and AR-Y445. Transformants were expanded in selective SC medium and spot plated on selective SC medium or selective SC medium supplemented with 5-FOA (1 g/L) for plasmid shuffle of plasmid 18 via *URA3* counter-selection^40^.

Fluctuation tests of the genomic *CAN1* gene were performed to determine genomic per-base substitution rates. To minimize propagation of *POL3* mutator strains, fluctuation tests were performed directly using colonies from plasmid shuffle plates. For each strain, 48 colonies from plasmid shuffle plates were individually scraped and resuspended in 120 μL 0.9% NaCl. 10 μL from each resuspension was diluted and subject to an event counts measurement via flow cytometry. This was done to identify colonies of similar cell count, because fluctuation tests are only appropriate when final population sizes for all replicates are similar. 24 resuspensions with similar event counts were used for fluctuation tests. 90 μL from the 24 resuspensions were mixed and plated onto SC medium lacking arginine and supplemented with 10X canavanine (0.6 g/L). 10 μL from four of the 24 resuspensions was pooled, diluted and titered on solid SC medium. Plates were incubated at 30 °C. Colonies were counted from titer plates after 2-4 days and from spot plates after 3-6 days. Based on titer counts, the average number of generations that occur during colony formation was ~15. Fluctuation data were analyzed as described above for *URA3* fluctuation data, but with mutation frequency parameters for *CAN1*^29^. The Fenton approximation^41^ was used to calculate the predicted rate of the extinct mutator strain.

To test the stability of mutator phenotypes, three colonies from each plasmid shuffle plate were passaged ten times at 1:100 dilutions (67 generations). A single clone from each final population was isolated and subject to *CAN1* fluctuation tests. This was performed using a protocol similar to that described previously for *URA3* fluctuation tests^23^, except cultures from *CAN1* fluctuation tests were plated onto solid SC medium lacking arginine and supplemented with 10X canavanine (0.6 g/L). Fluctuation data were analyzed as described above.

### Construction and characterization of p1-*Pf*DHFR strains

Strains GA-Y109, 149, 151 and 155 expressing *Pf*DHFR from p1 were derived from the parent strain, GA-Y102, via plasmid shuffle. GA-Y077 was constructed from AR-Y292 by concomitant deletion of genomically encoded *DFR1* and transformation of a centromeric plasmid encoding *Pf*DHFR. Pilot studies shown in **Extended Data Figure 5** used strains GA-Y077, 151 and 155. The results confirmed that strains dependent on *Pf*DHFR acquire sensitivity to pyrimethamine and evolve resistance by mutating p1-encoded *PfDHFR*.

### p1-*Pf*DHFR evolution experiment

GA-Y229, containing p1-encoded *Pf*DHFR replicated by TP-DNAP1-4-2, was serially passaged for large-scale evolution of pyrimethamine resistance. To start evolution, a saturated preculture of GA-Y229 was diluted 1:100 into 90 wells of a 96-well block containing 0.5 mL of selective SC supplemented with 100 μM pyrimethamine. (The remaining 6 wells served as controls, as described below.) The block was sealed and incubated at 30 °C. OD_600_ was monitored every 24 hours using a microplate reader (TECAN Infinite M200 PRO). When 80/90 experimental replicates reached an OD_600_ above 0.7, the entire block was passaged at a 1:100 dilution. If this cutoff was reached within 48 hours of the previous dilution, then the concentration of pyrimethamine was increased for all 90 replicates. Otherwise, pyrimethamine concentration remained the same. This was repeated until pyrimethamine concentration reached 3mM, the maximum concentration we were able to dissolve in SC media. The drug regimen proceeded from 100 μM to 500 μM, 1 mM, 2 mM, 2.5 mM and finally 3 mM. This regimen was guided by pilot experiments and designed to maintain strong selection throughout the experiment. Cultures were maintained at 100 μM for one passage, 500 μM for the second passage, 1mM for the third passage, 2 mM for passages 4-5, 2.5 mM for passages 6-7, and 3 mM for passages 8-13. During passage 4, the seal covering the 96-well block was punctured, so the passage was repeated from the passage 3 cultures stored at 4 °C. 50 μL volumes from each passage were stored with 25% glycerol at -80 °C.

Six randomly chosen control wells were filled with selective SC media lacking pyrimethamine, and two of these were seeded with GA-Y229. Media conditions in the control wells were kept the same throughout. No cross-contamination was detected and GA-Y229 grew robustly throughout.

After 80/90 replicates from passage 13 reached an OD_600_ of 0.7, the entire block was subject to whole-cell yeast DNA extraction. Bulk populations of p1 plasmids served as templates for PCR amplification of *PfDHFR*, and PCR amplicons were subject to Sanger sequencing. Mixed trace files were automatically annotated using Mutation Surveyor (SoftGenetics)^42^ and called mutations were manually verified. For focused analysis of the C50R, D54N, Y57H, C59R, C59Y and S108N mutations, trace file peak heights at bases 148, 160, 169, 175 and 323 in *PfDHFR* were converted to frequencies using QSVanalyzer^43^. The *PfDHFR* trace file of GA-Y229 served as the template of the homozygous w.t. base, at all five positions. Insertion frequencies at base 737 in *PfDHFR* were calculated from trace files using TIDE^44^ (default settings). Four *PfDHFR* mutations (Y70C, V125V, V146A, and V195A) were present at high frequency in GA-Y229 prior to pyrimethamine selection, after this strain was bottlenecked through a single cell during plasmid shuffle. These mutations were commonly observed as hitchhikers in evolved cultures, and were excluded from analysis.

Mutation frequencies calculated via QSVanalyzer^43^ were compared against frequencies calculated from deep sequencing. For deep sequencing analysis, *PfDHFR* was PCR amplified from DNA extract of two replicate populations. *PfDHFR* was amplified as two partially overlapping fragments. Amplicons were sent to Quintara Biosciences for library preparation and sequencing. At the sequencing vendor, the amplicons were additionally amplified to incorporate the TruSeq HT i5 and i7 adapters. The amplified libraries were sequenced on an Illumina MiSeq with the 500-cycle v2 reagent kit (Cat #: MS-102-2003). Paired-end reads were merged using PEAR^45^. Merged reads containing insertions or deletions were removed from analysis. Mutation frequencies match closely with results obtained from QSVanalyzer^43^. From one of the replicates, mutation frequencies calculated for the C50R, D54N, Y57H, C59R, C59Y and S108N mutations by QSVanalyzer^43^ are 2.3%, 60.5%, 66.3%, 95.5%, <1%, and 33.2%, respectively. In comparison, the corresponding frequencies from deep sequencing analysis are <0.1%, 57.1%, 54.7%, 97.3%, <0.1%, and 39.5%, respectively. From the other replicate, mutation frequencies calculated for the C50R, D54N, Y57H, C59R, C59Y and S108N mutations by QSVanalyzer^43^ are 5.9%, 35.8%, 51.7%, 97.4%, <1%, and 98.6%, respectively. In comparison, the corresponding frequencies from deep sequencing analysis are 2.5%, 34.1%, 52.9%, 99.6%, <0.1%, and 99.9%, respectively.

To track the dynamics of *Pf*DHFR evolution, mutational frequencies were tracked across all 13 passages of eight representative replicates. Cultures were inoculated from glycerol stocks into the same media condition they were last grown in. *PfDHFR* sequencing was performed as described for passage 13 of the large-scale evolution experiment. Mutation frequencies were calculated using QSVanalyzer^43^. For mutations that did not fully fix in any of the sequenced populations, a homozygous mutant allele was constructed by PCR and the resulting amplicon was similarly subject to Sanger sequencing.

### *Pf*DHFR MIC assay

The MIC of pyrimethamine was measured for 50 *Pf*DHFR alleles in the yeast strain, YH5^24^. Plasmids 31-80 in **Supplementary Table 7**, are yeast centromeric plasmids that express *Pf*DHFR from the weak *DFR1* promoter. Plasmids 31-80 were transformed into YH5^24^ and transformations were plated on selective SC medium supplemented with 100 μg/mL dTMP. After 5-6 days of growth at 30 °C, three transformants representing each allele were expanded in selective SC media supplemented with 100 μg/mL dTMP. Cultures were grown for 4 days at 30 °C. Saturated cultures were washed to remove dTMP. Resuspensions were diluted 1:100 into 50 μL volumes of 14 media conditions: YPD supplemented with 100 μg/mL dTMP, YPD, and YPD supplemented with 50 nM, 100 nM, 500 nM, 5 μM, 30 μM, 100 μM, 300 μM, 600 μM, 1 mM, 1.25 mM, 1.5 mM or 2 mM pyrimethamine. Inoculums were transferred into 384-well microplate reader trays, which were then sealed thoroughly to prevent evaporation and grown at 30 °C. Trays were unsealed, subject to OD_600_ measurement using a microplate reader (TECAN Infinite M200 PRO), resealed, and returned to the 30 °C shaker at 3-6 hour intervals for 7 days.

MIC was defined as log_10_ of the lowest pyrimethamine concentration (in nM) at which OD_600_ remained below 0.25 after 7 days of growth. MIC was individually calculated for three clones of the 50 *Pf*DHFR alleles. In total, 3 clones did not grow robustly in YPD supplemented with dTMP and were omitted from subsequent analysis. Of the 147 clones included in analysis, 6 clones did not exceed the MIC threshold at a low pyrimethamine concentration, but grew robustly at several higher concentrations. In these cases, we attribute failed growth to experimental error, and determined MIC as if growth were sustained in the aberrant condition.

### p1 copy number control

To titrate p1 copy number, the catalytically inactive TP-DNAP1 (D641A) variant was placed under the control of the repressible *MET3* promoter in AR-Y062, which expresses w.t. TP-DNAP1 and the mKate2 reporter from p1. Strains were grown in SC media containing methionine, ranging in concentration from 0 μM to 450 μM, for 2 days at 30 °C. Afterwards, mKate2 fluorescence was measured, as described above, to assay p1 copy number.

---

**Extended Data Figure 1.**
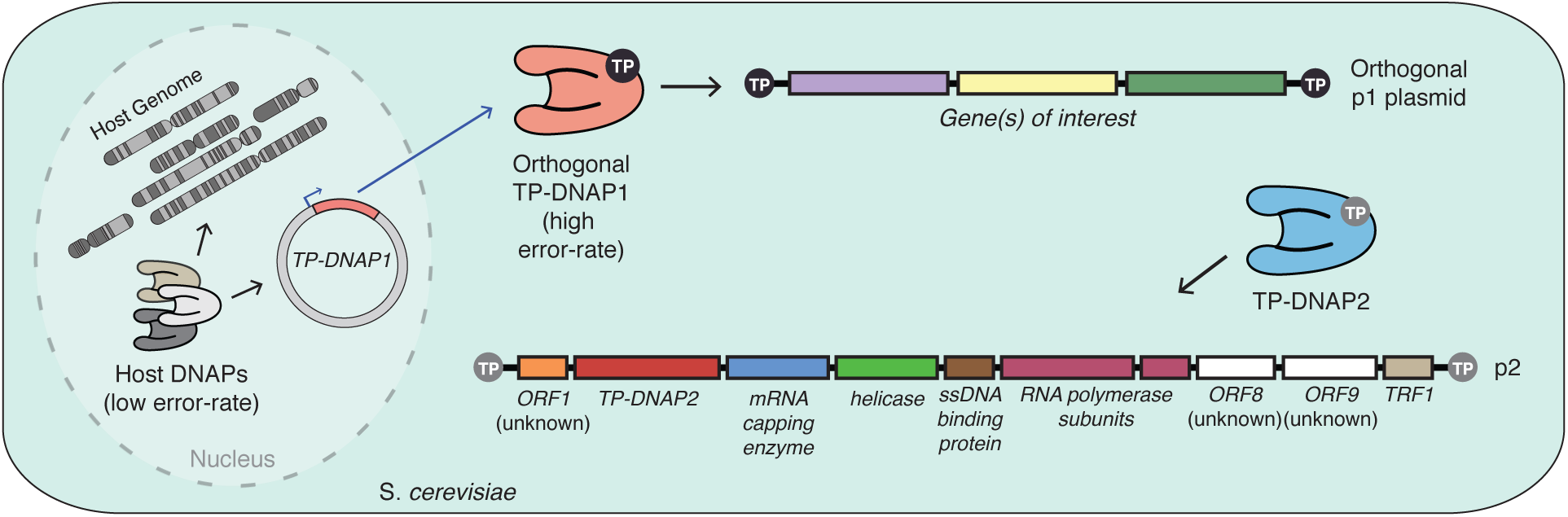
An expanded view of OrthoRep. The specific basis for OrthoRep is the p1/2 (also known as the pGKL1/2) plasmid system. p1 and p2 are linear, high-copy, double-stranded DNA plasmids that propagate autonomously in the cytoplasm of *S. cerevisiae*. TP-DNAP1 expressed from a nuclear plasmid replicates p1. TP-DNAP2 expressed from p2 replicates p2. TP-DNAP1 and TP-DNAP2 use terminal proteins (TPs) covalently attached at the 5’ ends of p1 and p2, respectively, as replication origins for TP-primed replication. All of the accessory components required for replication and transcription are encoded on p2. TP-DNAP1 does not replicate p2 and TP-DNAP2 does not replicate p1 (manuscript in preparation), meaning that the high error rate of p1 replication does not affect p2-encoded genes and the low error-rate of p2 replication does not compete with p1 mutagenesis. ORFs with unknown function are indicated. TRF, Terminal Recognition Factor.

**Extended Data Figure 2.**
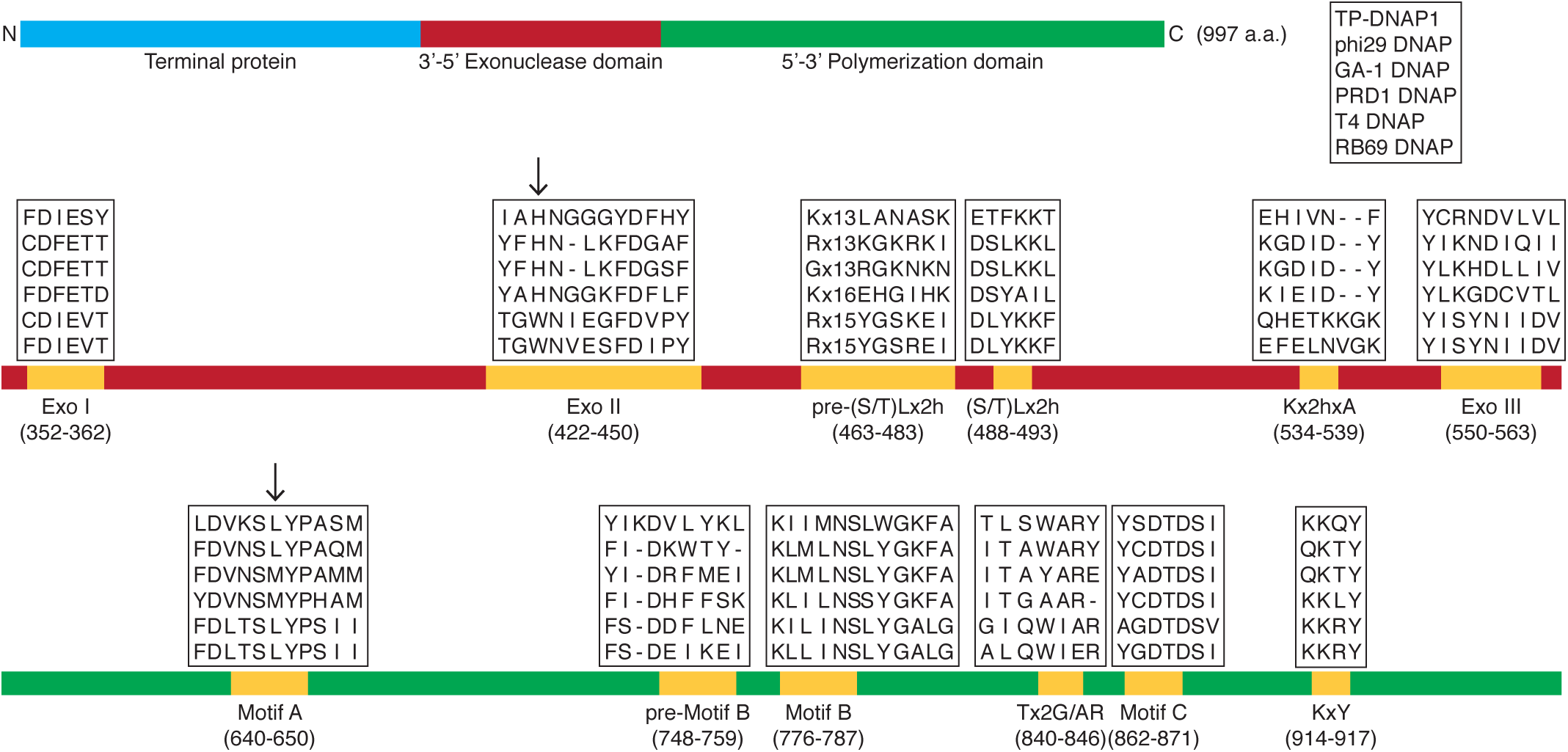
Homology analysis of TP-DNAP1 fidelity regions. The architecture of TP-DNAP1 consists of a fusion between the terminal protein, a 3’-5’ proofreading exonuclease domain, and a DNA polymerization domain. Motifs responsible for fidelity in the exonuclease and proofreading domains are highlighted. A multiple sequence alignment between TP-DNAP1 and five closely related family B DNAPs is shown. In the larger homology study described in the main text, multiple sequence alignment with 99 closely related DNAPs (**Supplementary Table 1**) was used to identify positions that exhibit amino acid (a.a.) variation and are flanked by conserved residues (**Methods**). Two candidate positions identified from this study are denoted with arrows. Amino acid variations found at these positions were transplanted into the corresponding location in TP-DNAP1. A total of 87 such TP-DNAP1 mutants were generated and screened in OR-24. Twenty-four of the TP-DNAP1 variants displayed elevated mutation rates, but almost 60% of these suffered from low activity, judging by the copy number of p1 (**Supplementary Table 2**).

**Extended Data Figure 3.**
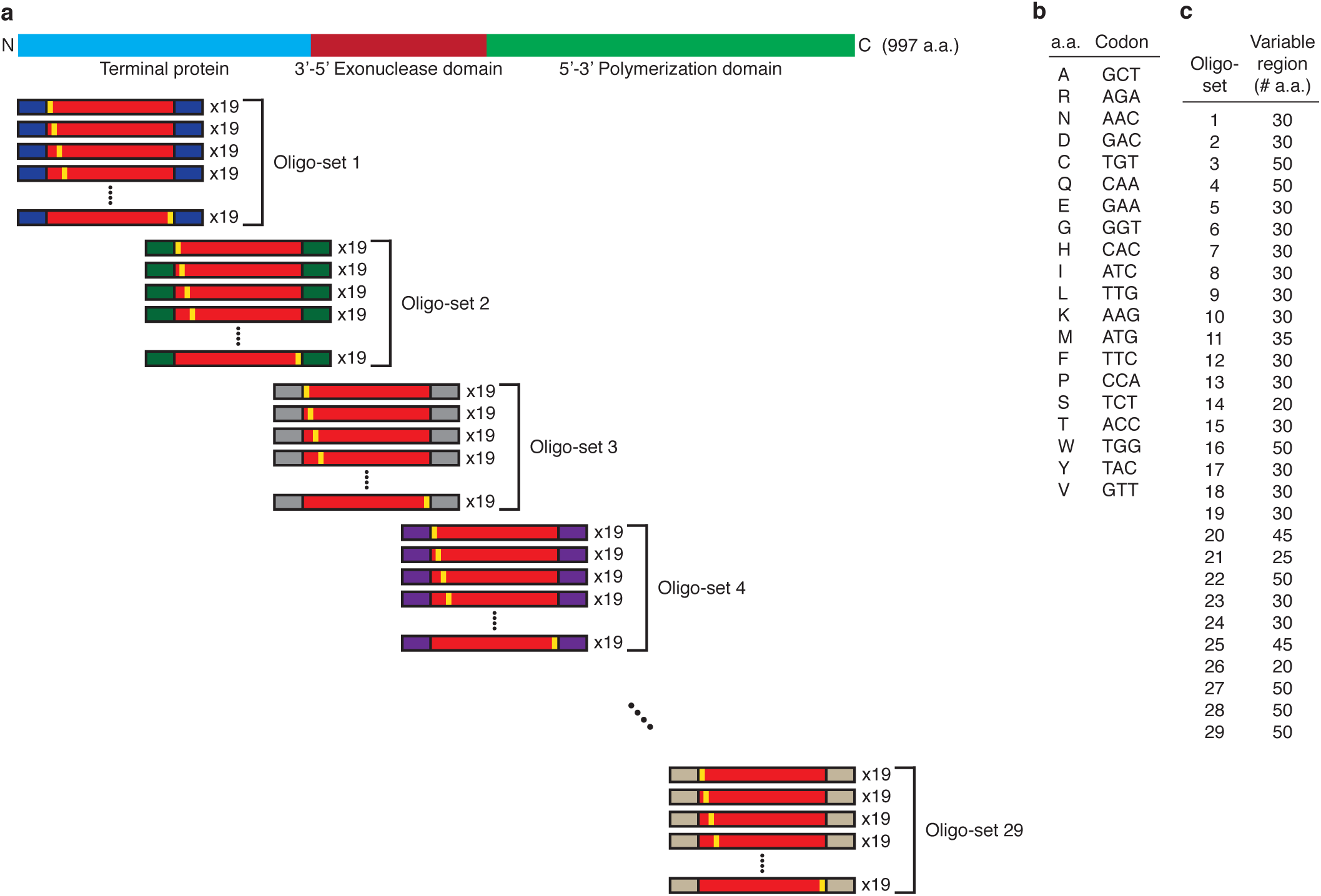
Design of a TP-DNAP1 scanning saturation mutagenesis library. **a.**, A pool of ~19,000 oligonucleotides ranging in length from 130-200 nt were designed as 30 sets, each encoding a 20-50 amino acid (a.a.) variable region flanked by ~25 bp constant regions. Variable regions mutate each w.t. codon to 19 codons, representing all single amino acid substitutions. Each oligo set was PCR amplified and assembled with corresponding TP-DNAP1 plasmid backbones, yielding 30 full-length TP-DNP1 plasmid sub-libraries. **b**, The genetic code used for mutagenesis, chosen to maximize the codon adaptation index in *S. cerevisiae*. **c**, Lengths of variable regions from each oligo set. Oligo sets 3 and 4, which were synthesized separately from the rest, have overlapping variable regions. Oligo sets 1-10 correspond to the putative TP portion of TP-DNAP1.

**Extended Data Figure 4.**
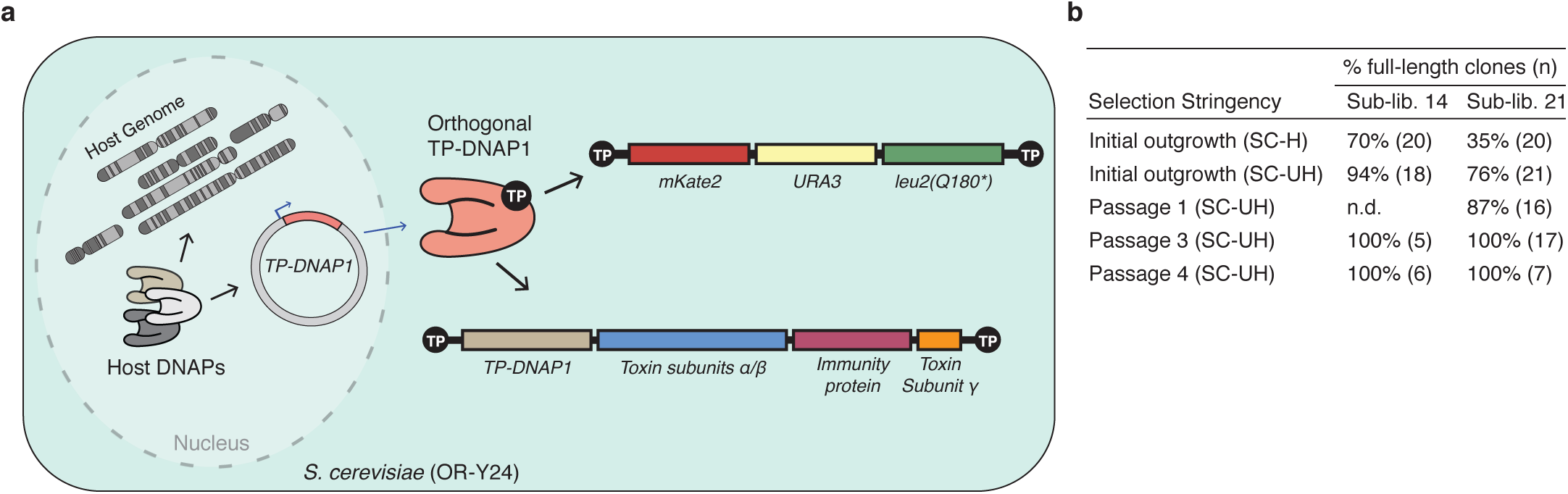
Functional purification of TP-DNAP1 sub-libraries. **a**, A conceptual illustration of OR-Y24, which is a meta-stable strain containing a mixture of w.t. p1 and recombinant p1 encoding *mKate2*, *URA3*, and *leu2(Q180*)*. Under selection for URA3 in media lacking uracil, recombinant p1 increases in copy number and displaces w.t. p1 because they share the same source of TP-DNAP1. Loss of w.t. p1 decreases the amount of TP-DNAP1 available to both plasmids, which initiates a feedback loop that drops the copy number of both plasmids and drives OR-Y24 to extinction. Functional TP-DNAP1 variants expressed *in trans* from a nuclear *CEN6/ARS4* plasmid can replicate recombinant p1 at a constant level and thereby rescue growth. **b**, Functional purification of representative TP-DNAP1 sub-libraries 14 and 21. OR-Y24 cells transformed with sub-libraries 14 or 21 were split and inoculated into outgrowth conditions that were selective only for plasmid uptake (SC-H) or for plasmid uptake and *URA3* expression (SC-UH). Selection in SC-UH was maintained for four 1:100 serial passages. After each passage, TP-DNAP1 plasmids were isolated from individual clones and subject to Sanger sequencing. Data shown are the percentage of (n) clones encoding full-length TP-DNAP1 variants.

**Extended Data Figure 5.**
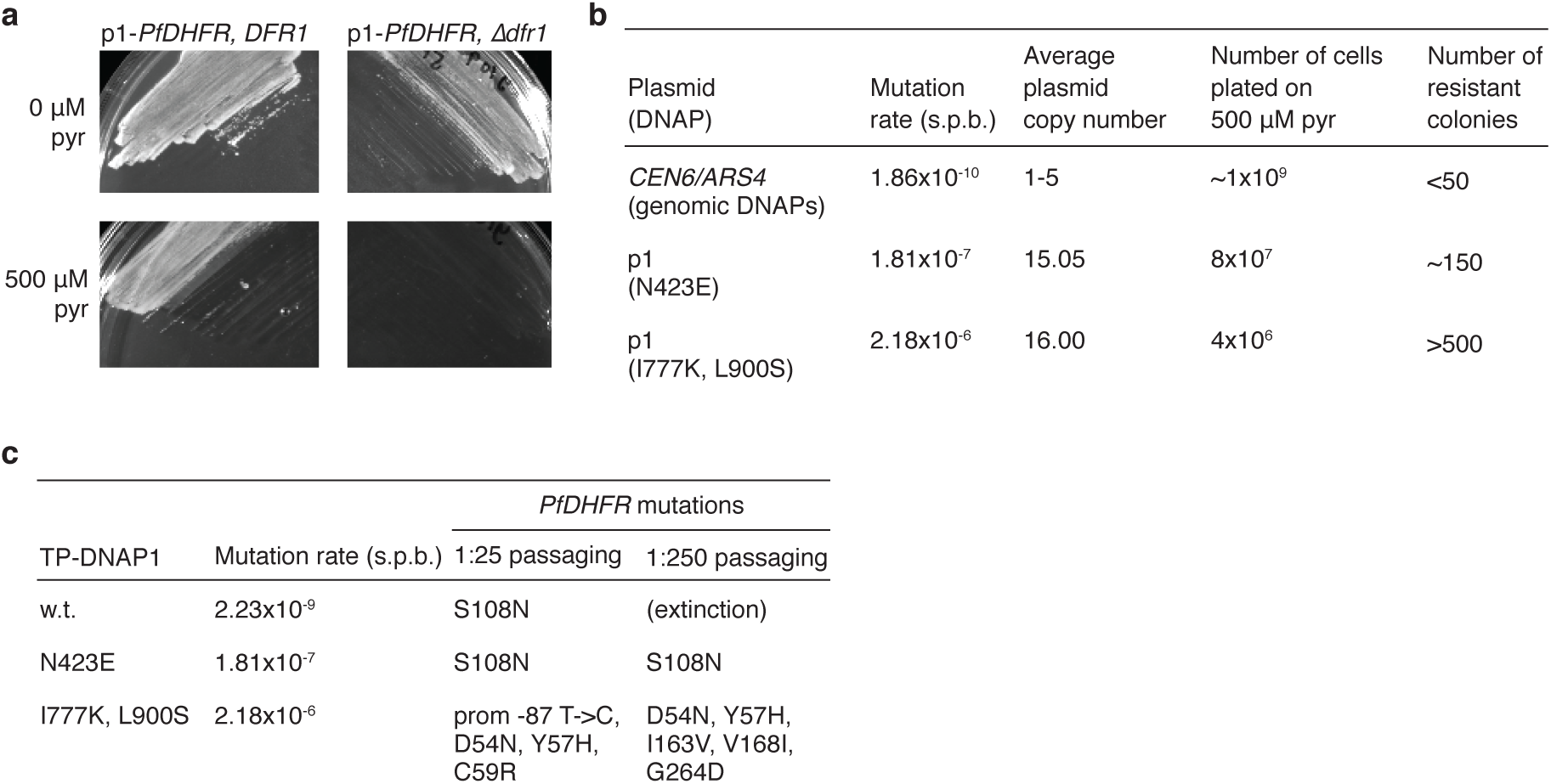
Pilot studies of p1-PfDHFR evolution. **a**, Yeast strains dependent on p1-encoded PfDHFR acquired sensitivity to pyrimethamine (pyr). *Pf*DHFR was expressed from p1 in strains that retain or lack *DFR1*, which encodes yeast’s endogenous DHFR. Strains were grown in selective SC media and plated on solid media with or without 500 μM pyrimethamine. Plates were incubated at 30 °C for 5 days prior to imaging. **b**, Pyrimethamine resistant clones arise in small culture volumes. A yeast strain that encodes *PfDHFR* on a nuclear plasmid and two OrthoRep strains that encode PfDHFR on p1 at for rapid mutation were grown to saturation in selective SC media and plated on solid media supplemented with pyrimethamine. After 5-6 days of growth at 30 °C, resistant colonies were counted. p1-encoded *PfDHFR*s carried resistance mutations in all 30 resistant clones sequenced. **c**, OrthoRep strains evolved pyrimethamine resistance in batch culture by rapidly mutating p1-encoded *PfDHFR*. OrthoRep strains with varying p1 mutation rates were serially passaged in 25 mL cultures, at 1:25 or 1:250 dilutions, in selective SC media initially supplemented with 500 μM pyrimethamine. OD_600_ was monitored daily and saturated cultures were passaged into gradually increasing drug concentrations as cultures adapted. After strains evolved resistance to 2 mM pyrimethamine, bulk populations of p1 plasmids were extracted and subject to Sanger sequencing. The OrthoRep strain containing w.t. TP-DNAP1 stopped growing in the 500 μM pyrimethamine condition when passaged at 1:250 dilutions.

**Extended Data Figure 6.**
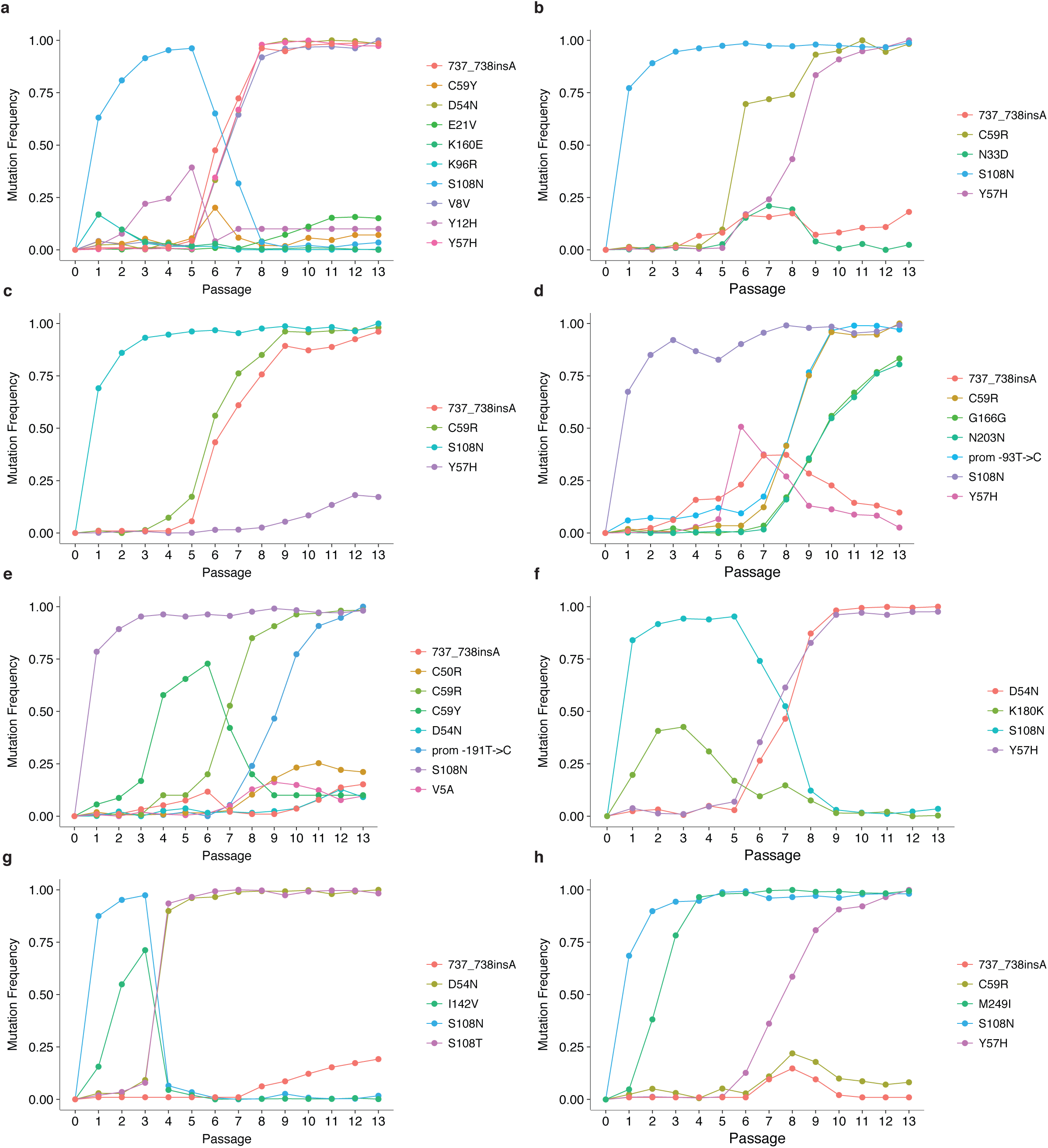
Dynamics of PfDHFR evolution in eight representative populations. Mutation frequencies were tracked across all 13 passages. Populations from each passage were revived from glycerol stocks in the same media condition that they were initially grown in. Mutation frequencies were calculated from Sanger sequencing of revived populations (see **Methods** for details of SNP analysis).

**Extended Data Figure 7.**
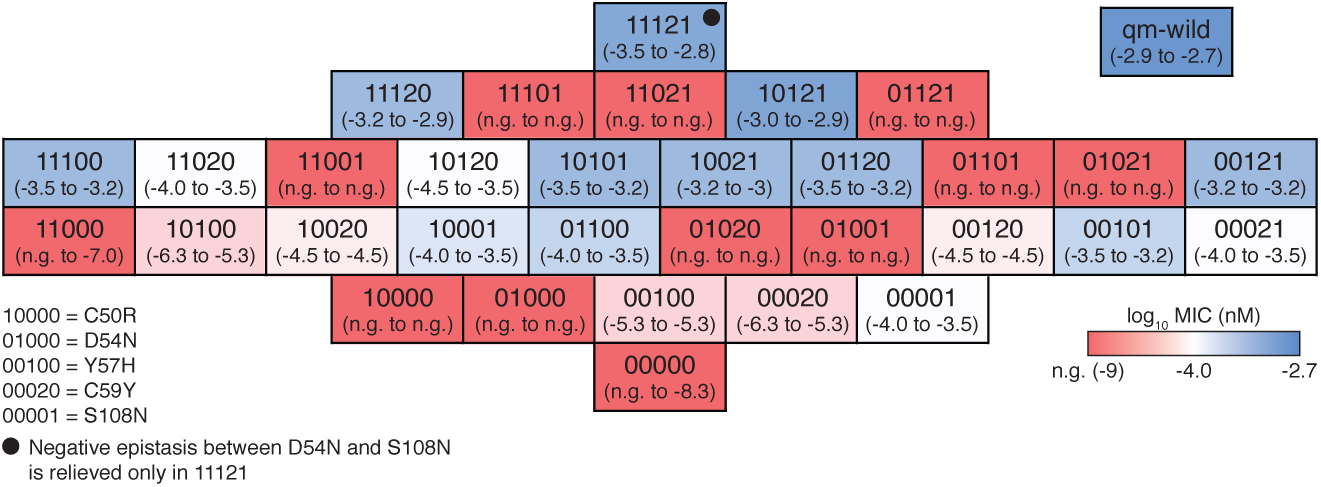
A fitness map of a five-mutation PfDHFR landscape defined by C50R, D54N, Y57H, C59Y and S108N. Seven out of 78 adapted populations contain high frequencies of C59Y (replicates 1, 7, 38, 39, 41, 61, and 76 in **Figure 3B**) and access an alternative five-mutation landscape. MIC of pyrimethamine was determined for yeast strains expressing all 32 *PfDHFR* alleles from this landscape (**Methods**). Data shown are the range of log_10_(MIC of pyrimethamine (nM)) for biological triplicates, with a color on a red-blue scale indicating the median. The mid-point of the red-blue scale is shifted to distinguish highly resistant alleles. n.g., no growth.

**Extended Data Figure 8.**
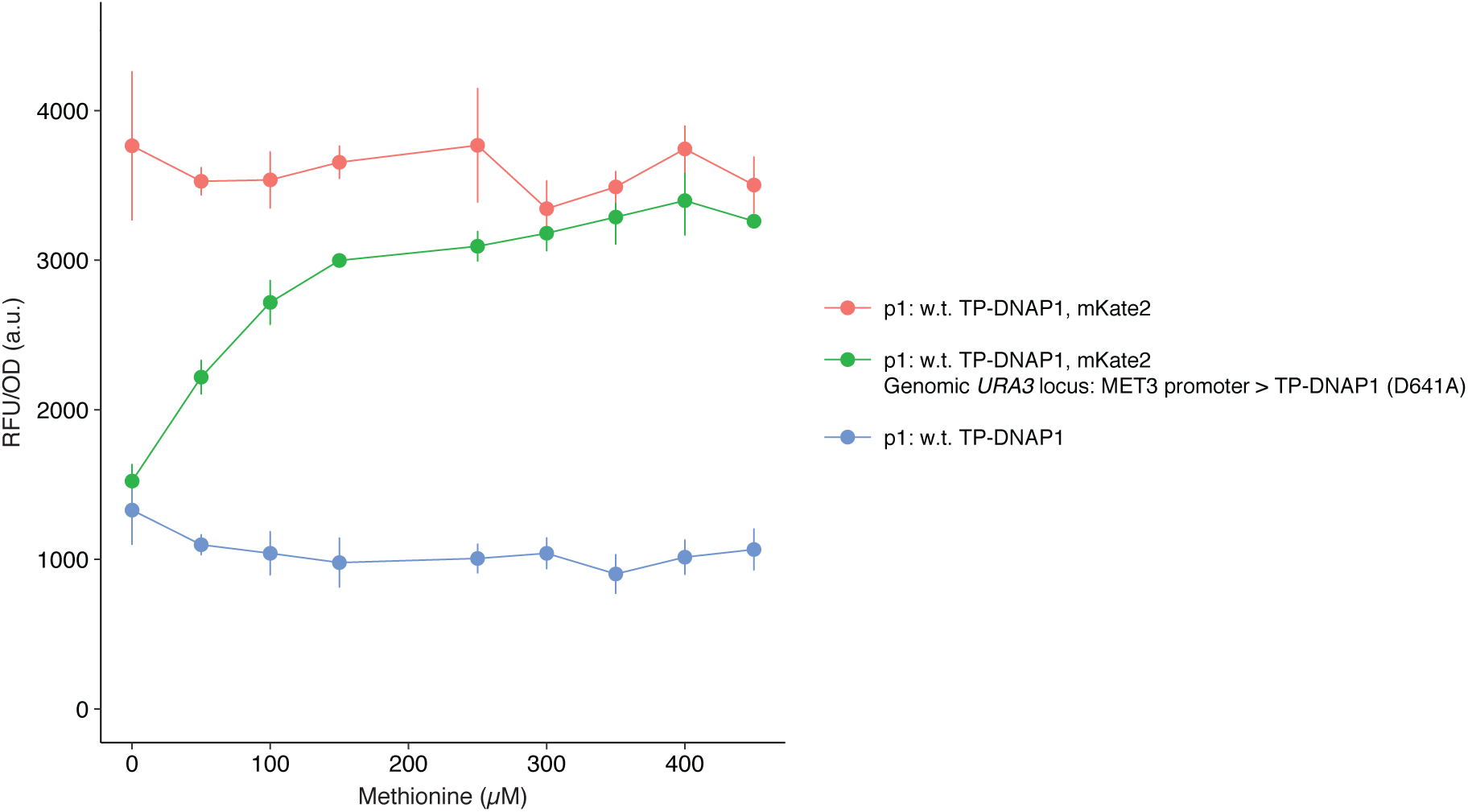
Titratable control of p1 copy number. TP-DNAP1 encodes a highly conserved catalytic residue at D641. The TP-DNAP1 (D641A) variant is unable to polymerize DNA, but can still compete with fully functional TP-DNAP1 for replication initiation at the TP origin. TP-DNAP1 (D641A) was expressed under the control of the repressible *MET3* promoter in a strain expressing w.t. TP-DNAP1 and mKate2 from p1. Strains were grown in SC media supplemented with methionine for two days, after which OD_600_ and mKate2 fluorescence were measured. Data shown are the mean OD_600_-normalized mKate2 fluorescence ± standard deviation (measured in arbitrary units (a.u.)) for biological triplicates.

**Extended Data Table 1.**
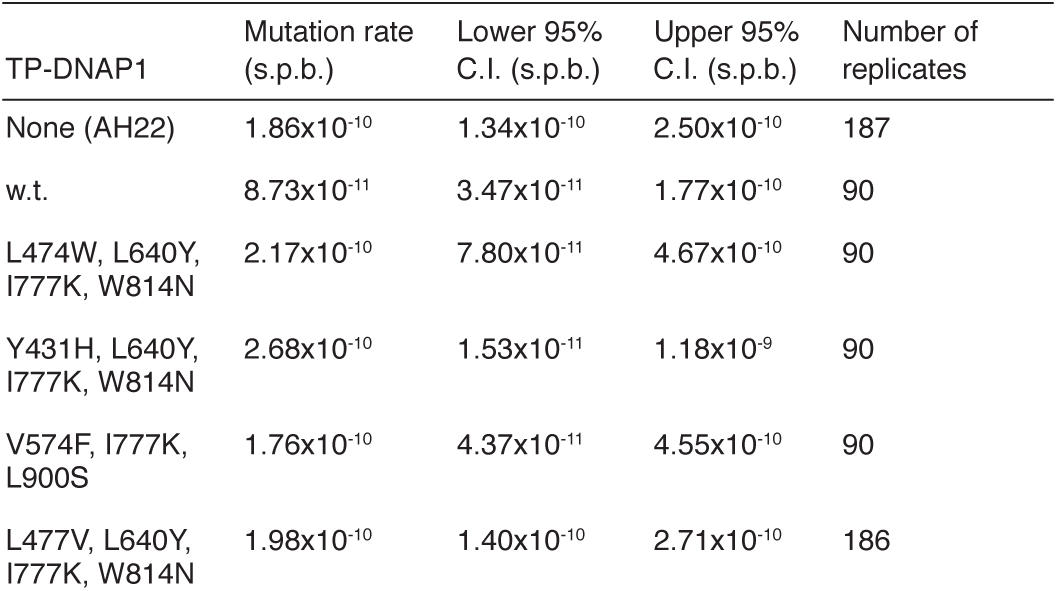
Yeast genomic substitution mutation rates in the presence of OrthoRep. Per-base substitution rates were measured at the *URA3* locus in the presence of TP-DNAP1 variants. Genomic fluctuation tests were performed in the presence of p1 replication by TP-DNAP1 variants by selecting for a p1-encoded marker. AH22 is the parent OrthoRep strain and lacks p1 and TP-DNAP1. Data shown are individual mutation rate measurements, each with a corresponding 95% confidence interval (C.I.) and a count of the number of replicates performed for the fluctuation test.

**Extended Data Table 2.**
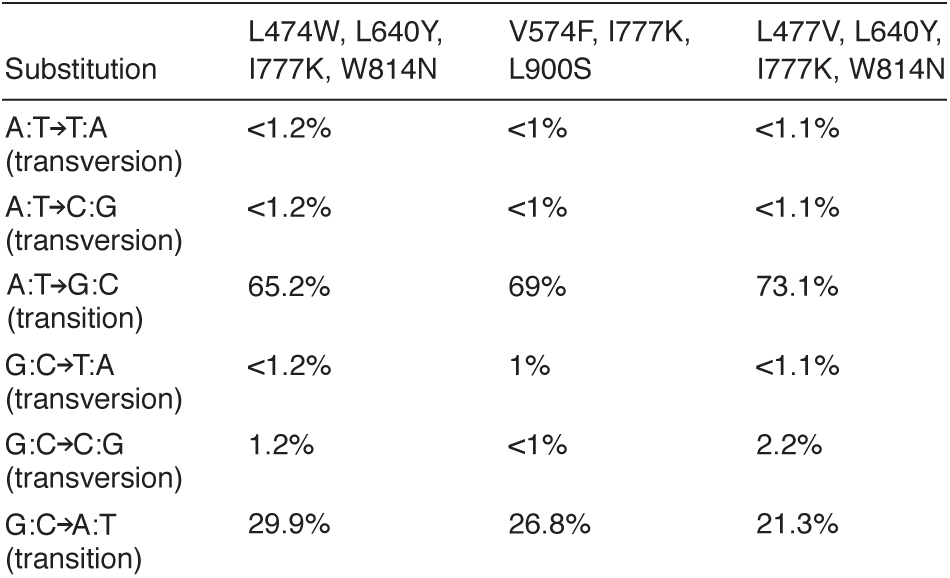
Substitution mutation preferences of highly error-prone TP-DNAP1s. All substitution preferences except for the G:C A:T transition were measured from reversion of *leu2 (538C>T, 540A>G)*. The substitution rate of G:C A:T was determined from fluctuation tests of *ura3 (278A>G)*. The per-base-normalized G:C A:T substitution rates of TP-DNAP1 (L474W, L640Y, I777K, W814N), TP-DNAP1 (V574F, I777K, L900S), and TP-DNAP1 (L477V, L640Y, I777K, W814N) are 6.41×10^−6^ s.p.b, 1.73 ×10^−5^ s.p.b, and 1.77 ×10^−5^ s.p.b, respectively. These rates were incorporated in proportion to the individual error rates calculated for each of the other five substitutions (see **Methods** for details). Data shown are normalized percentages of each substitution mutation.

**Supplementary Table 1.**
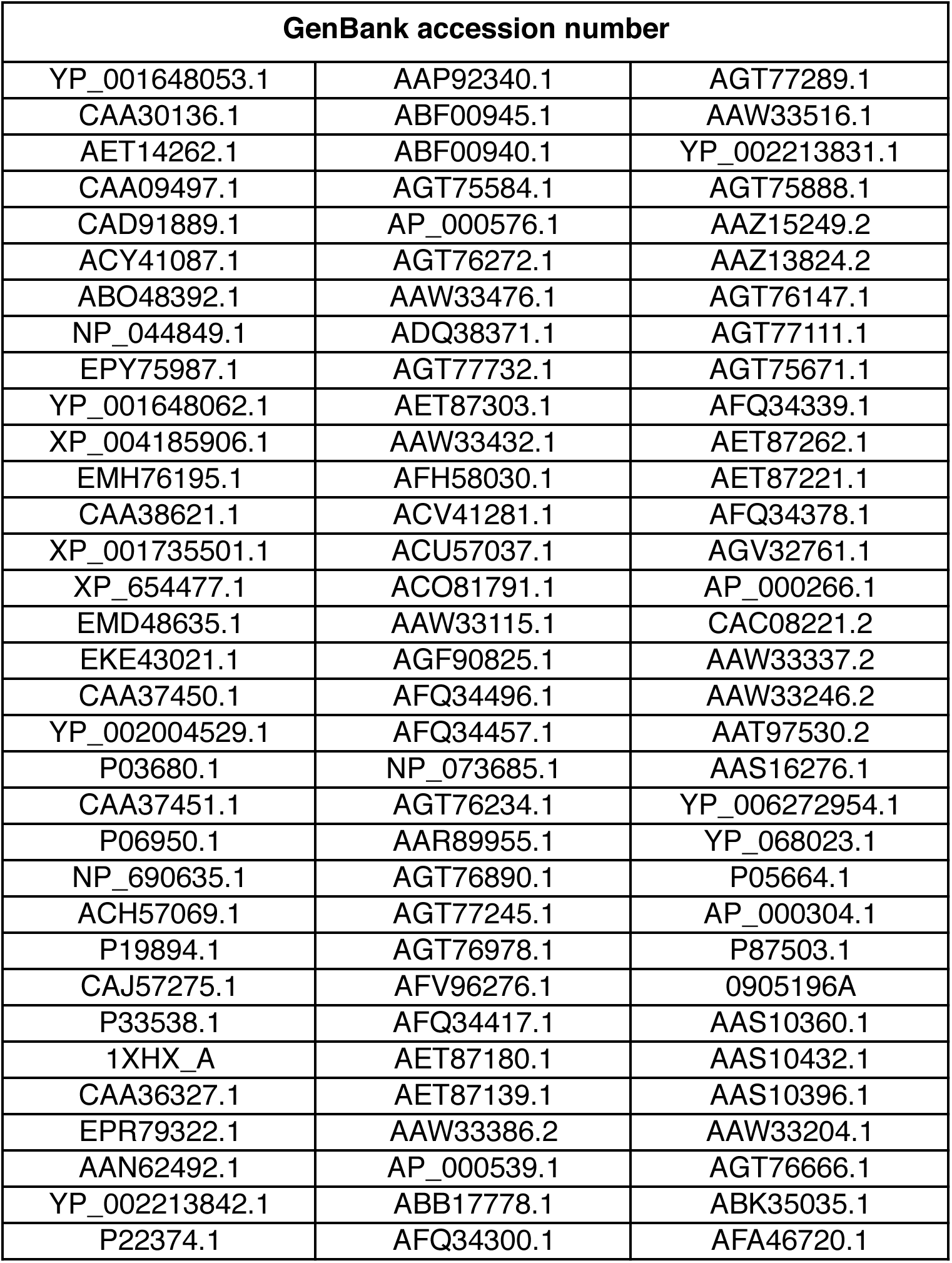
99 TP-DNAP1 homologs generated via protein BLAST^34^.

**Supplementary Table 2.**
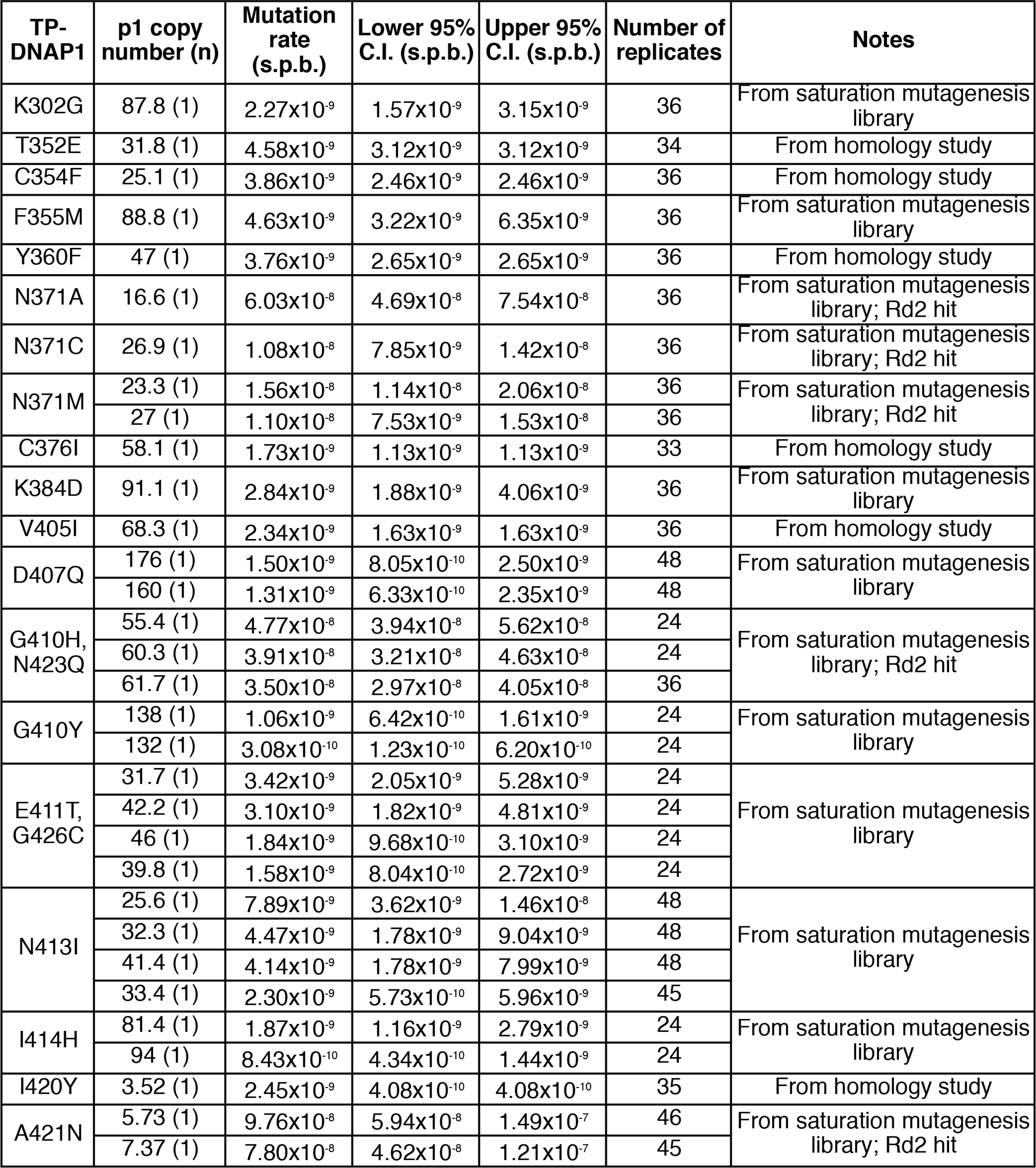
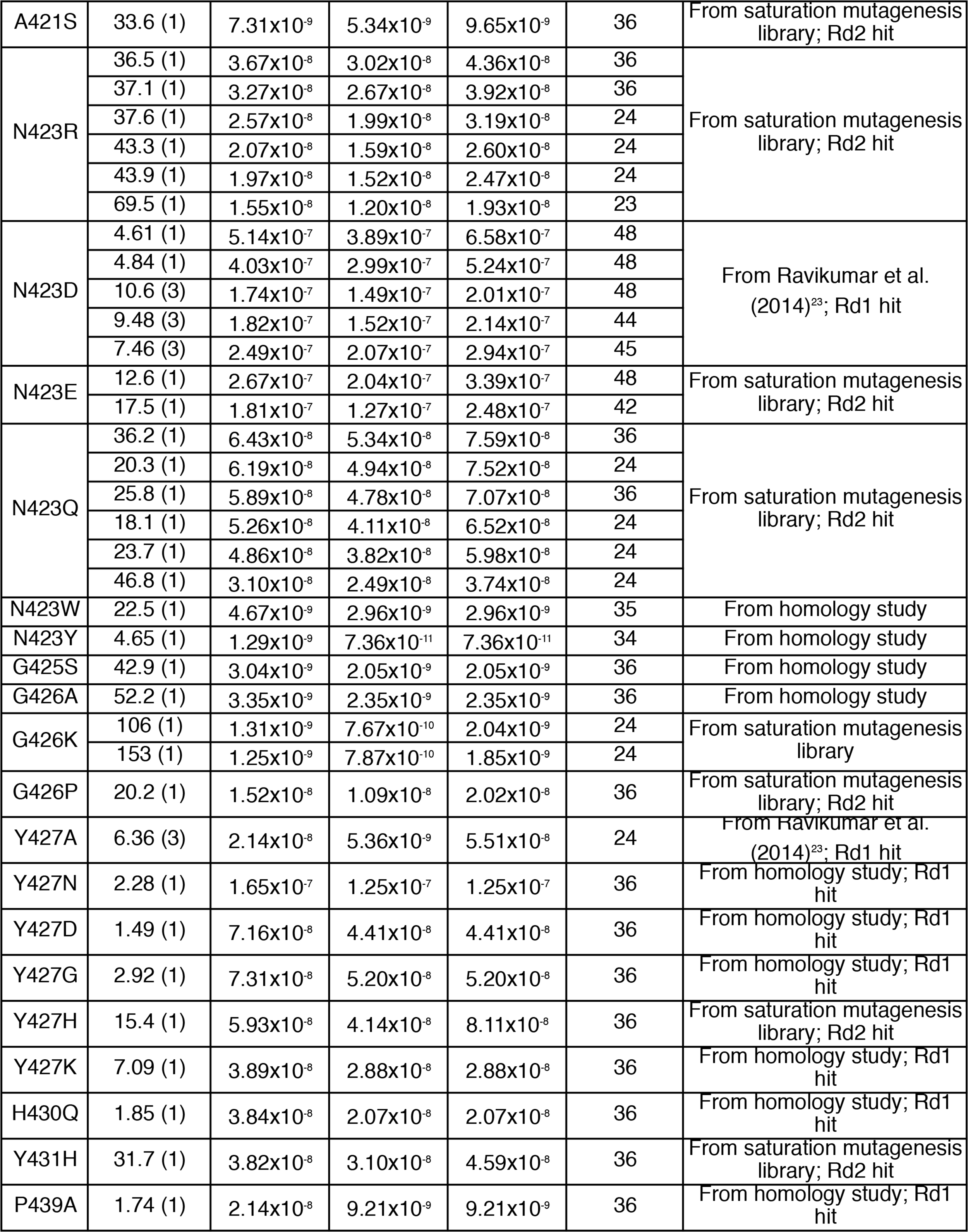
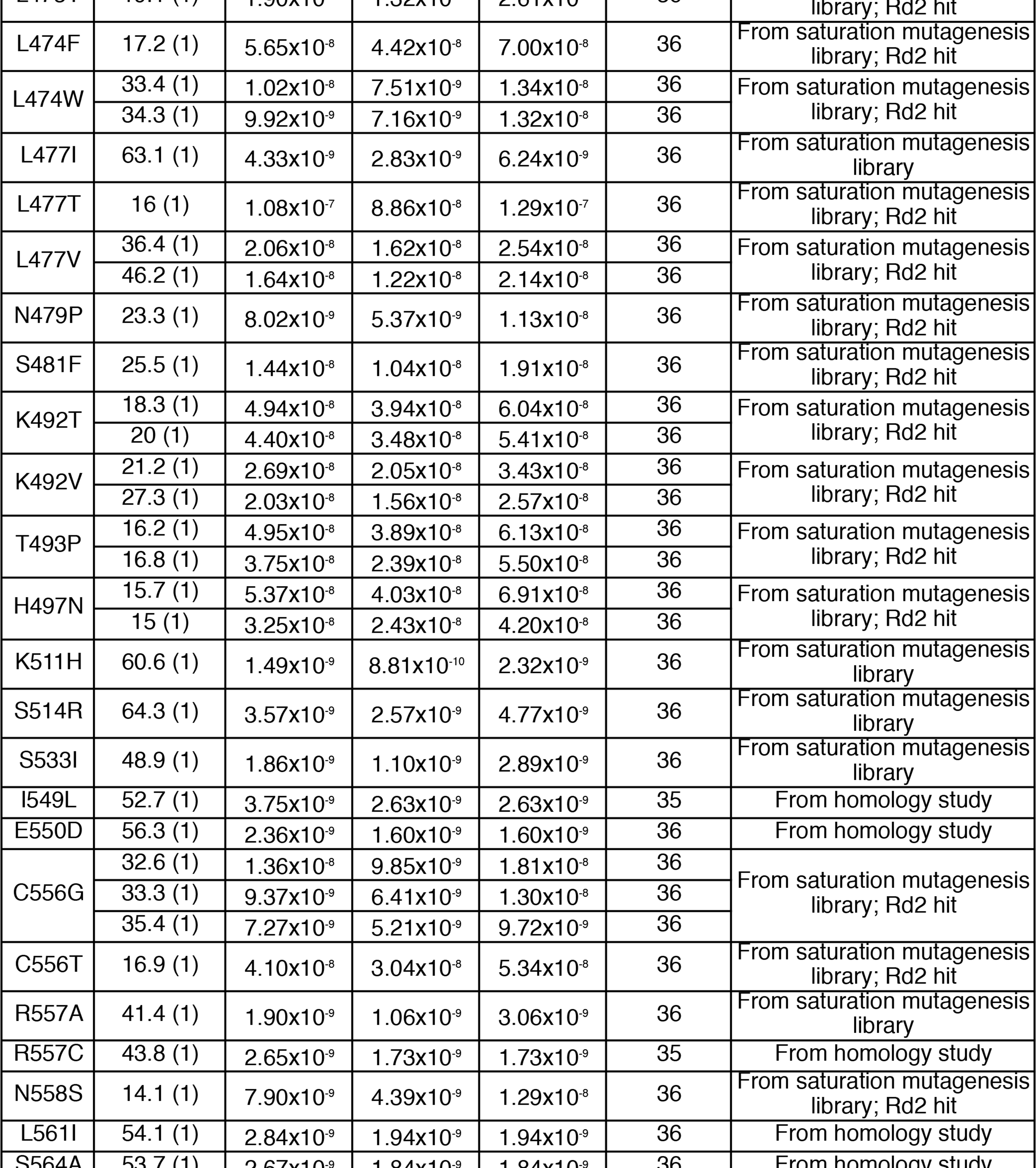
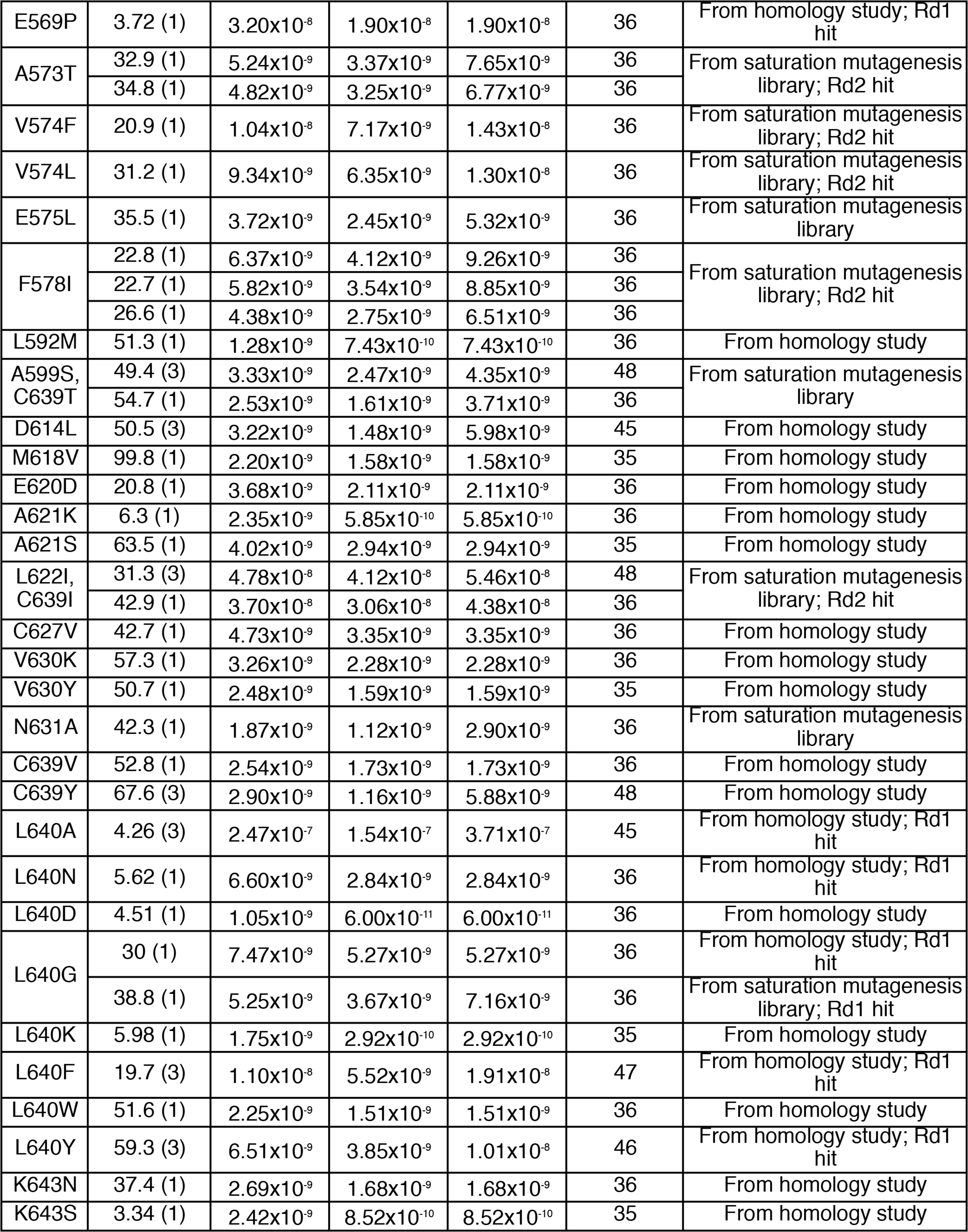
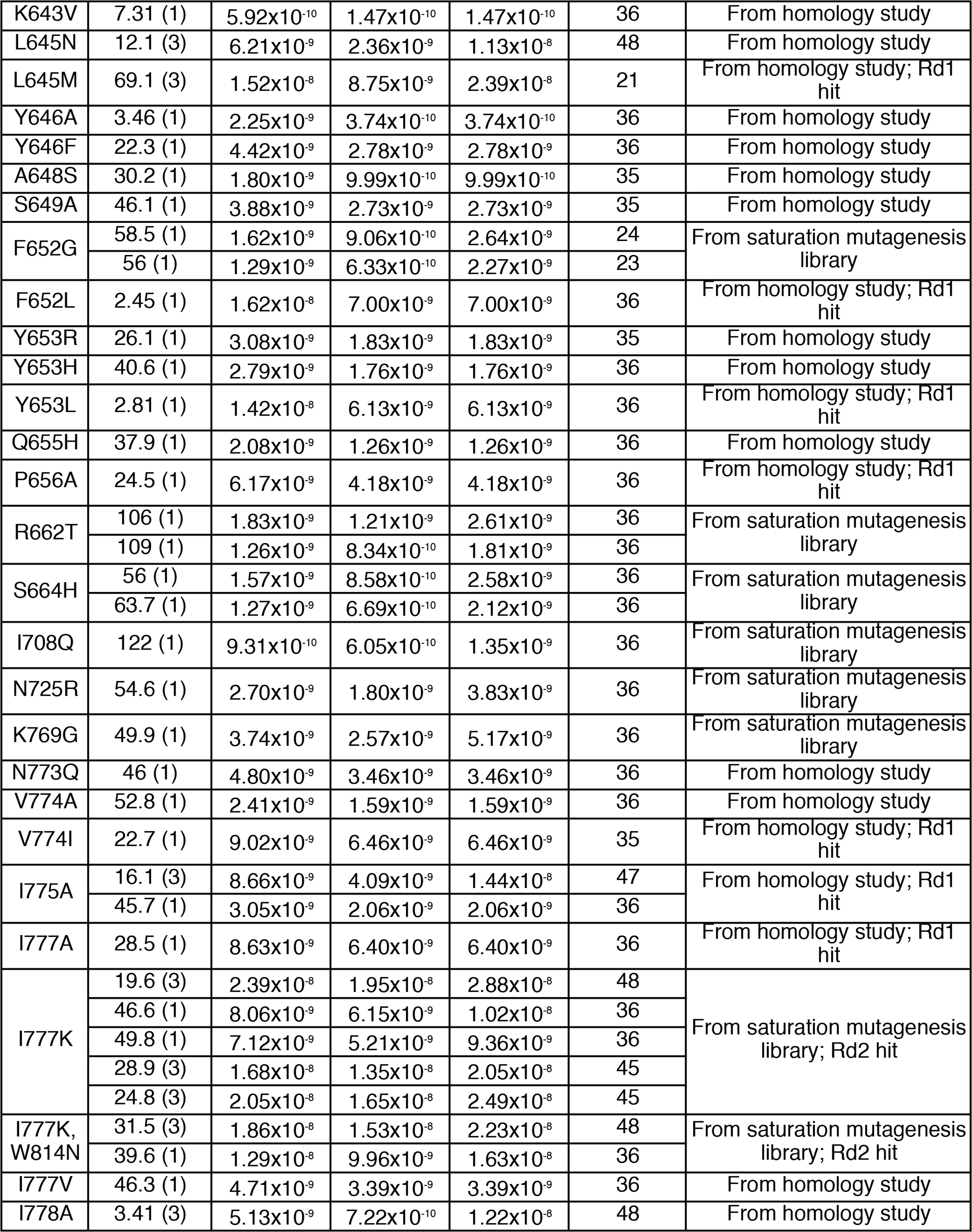
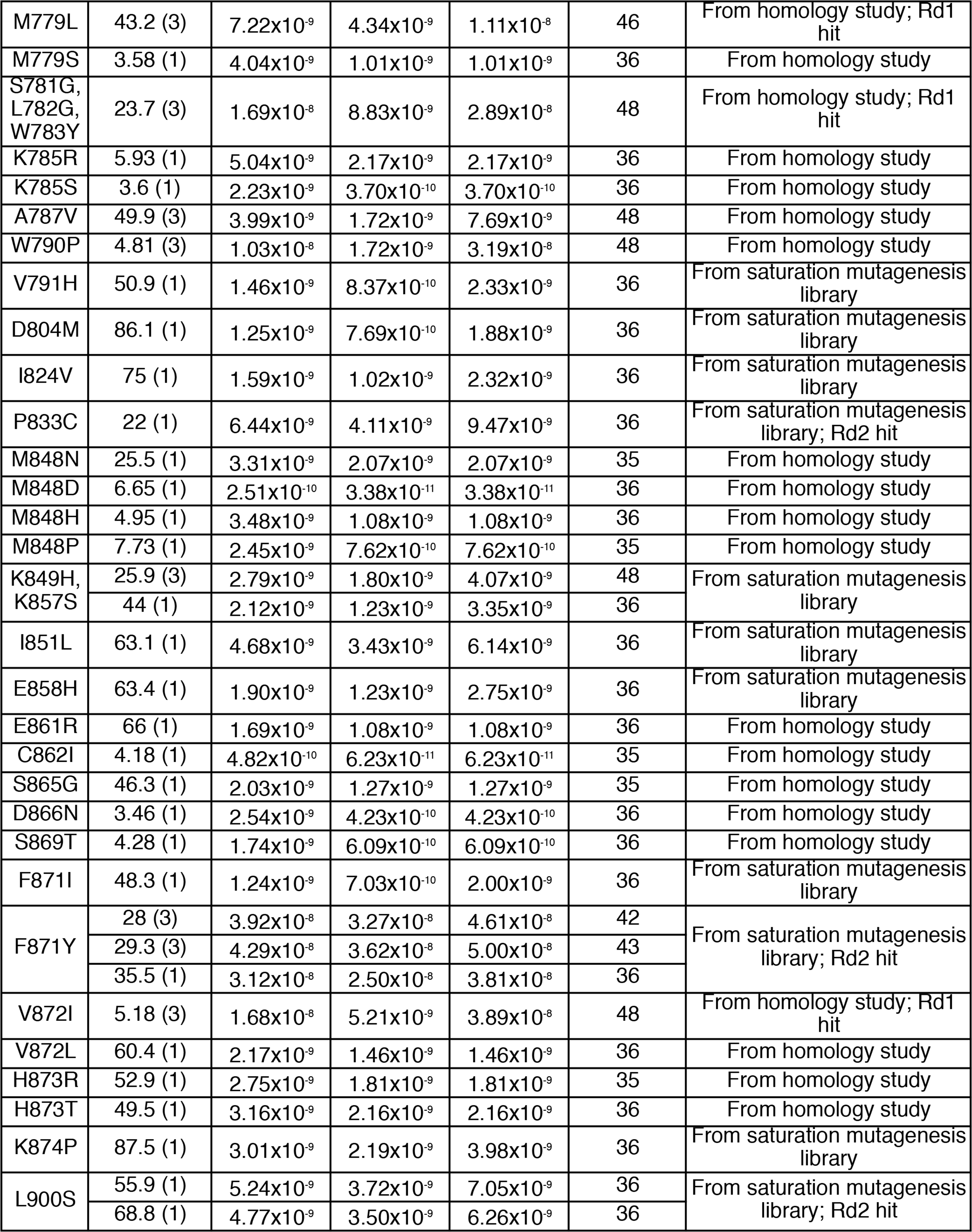
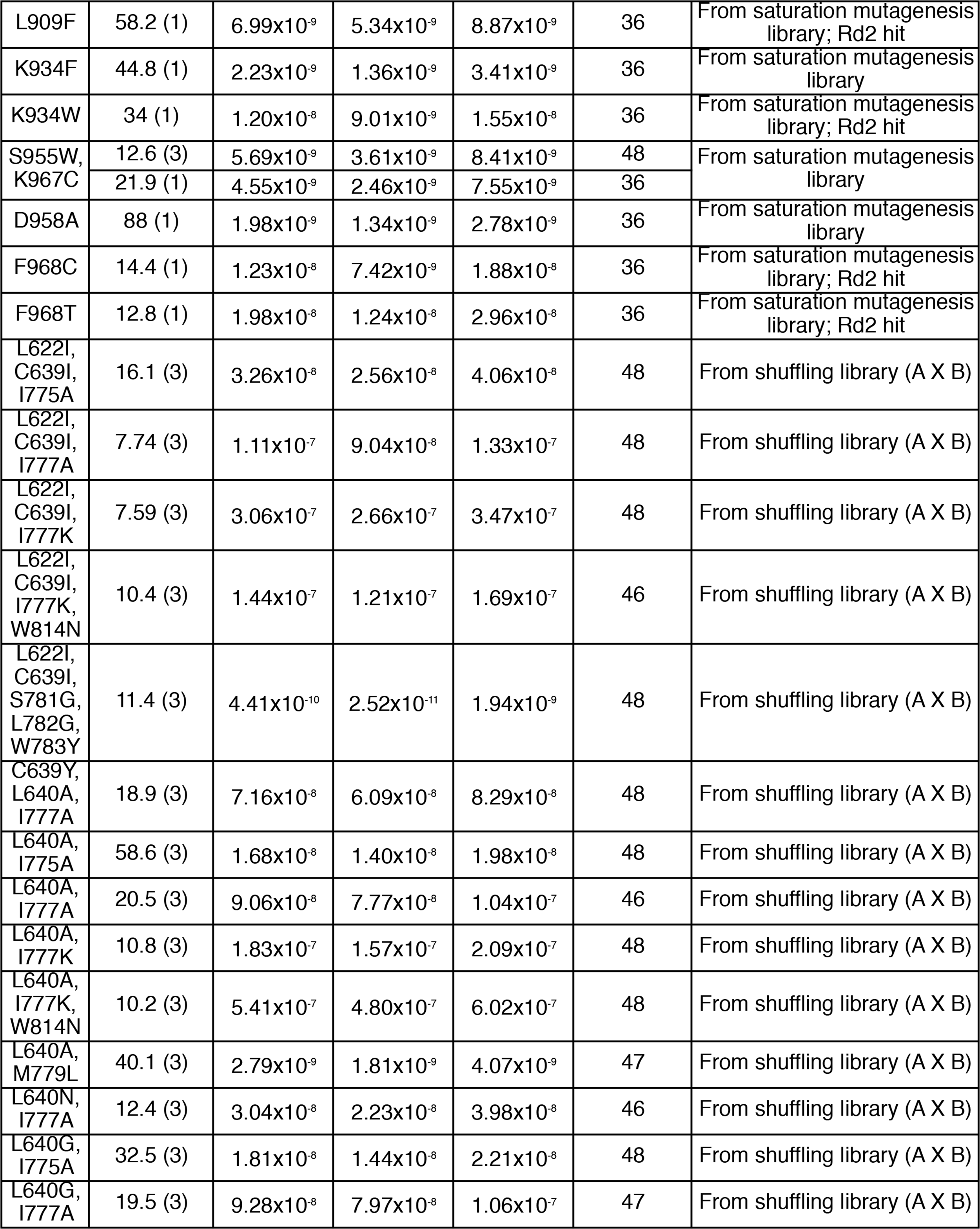
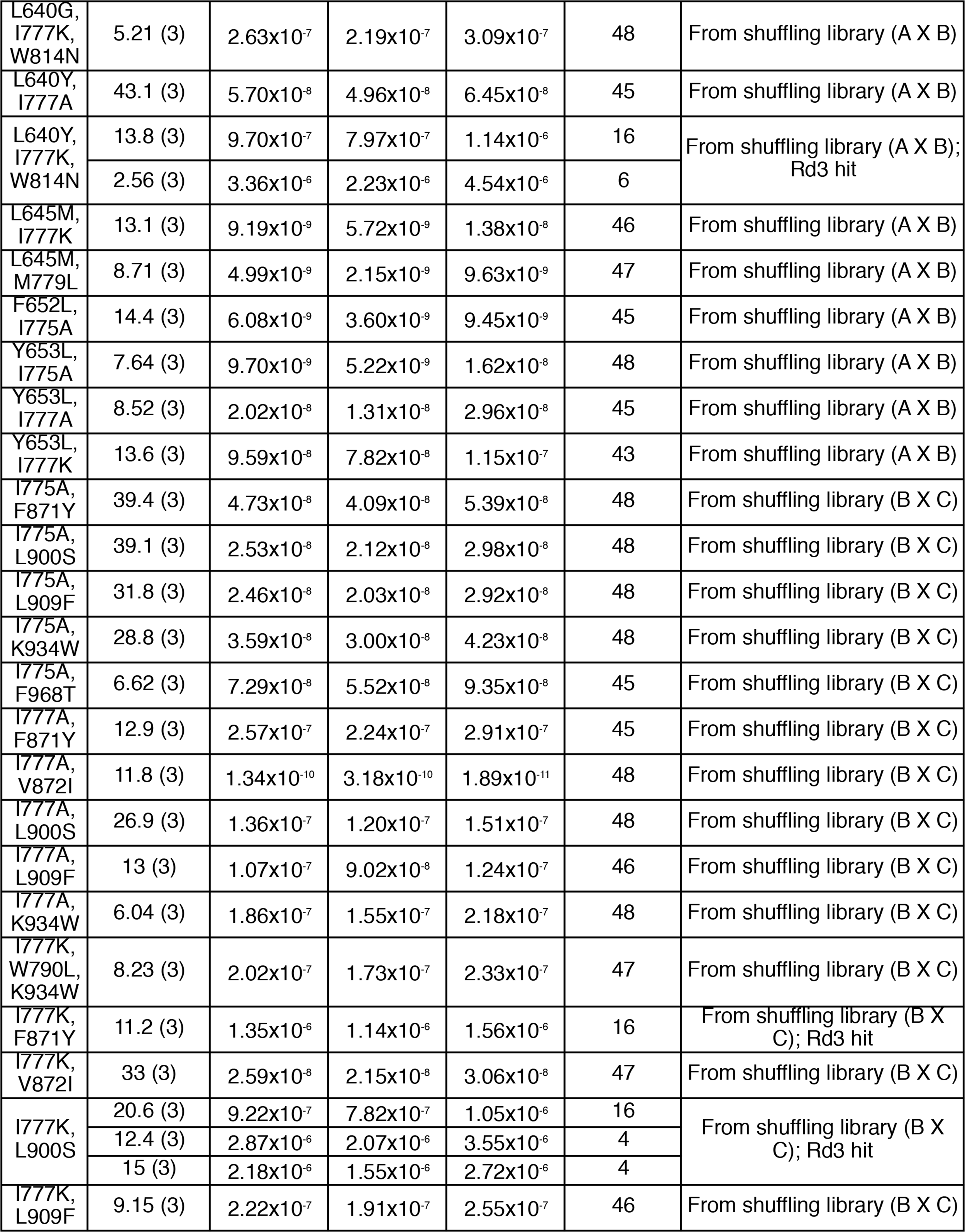
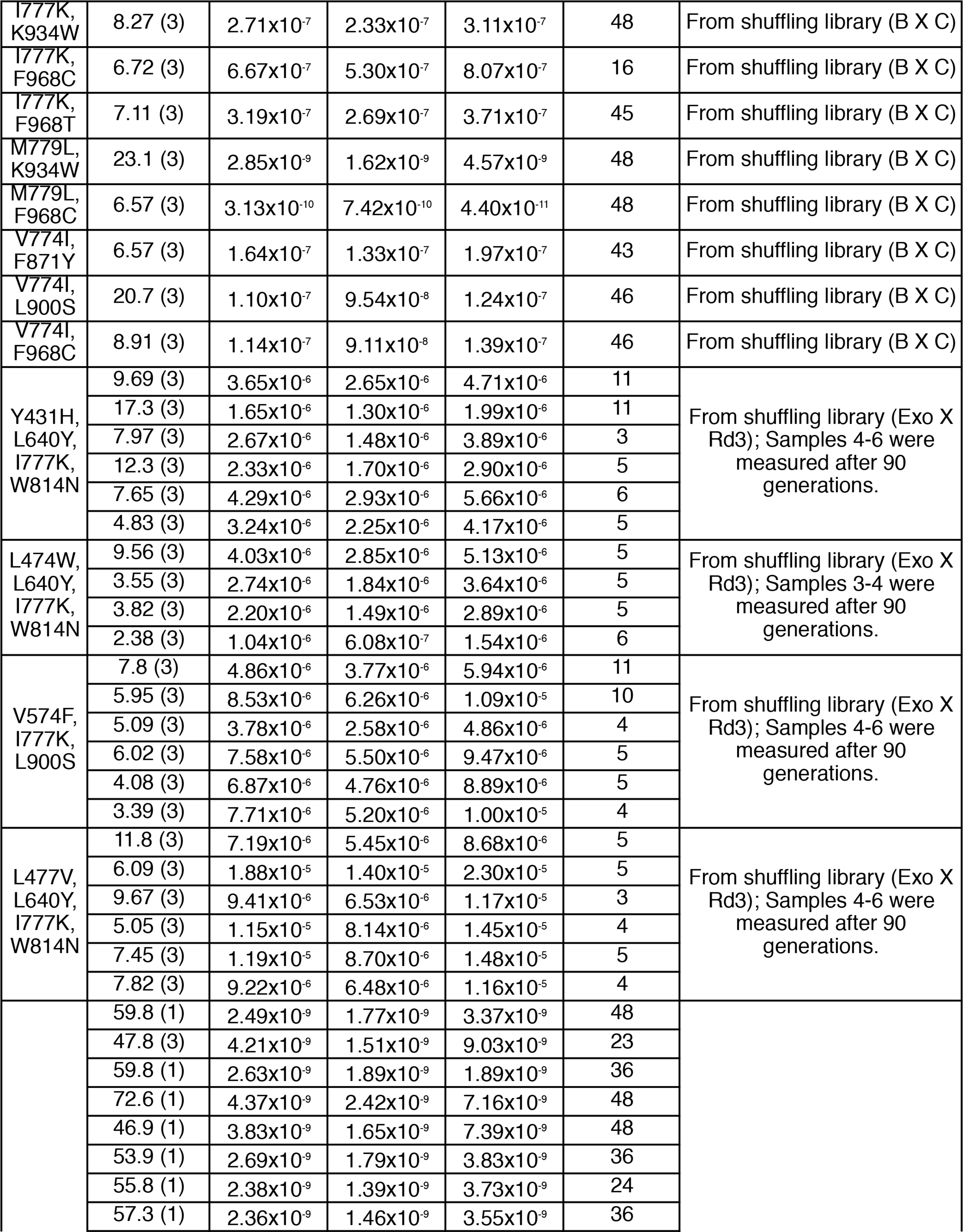
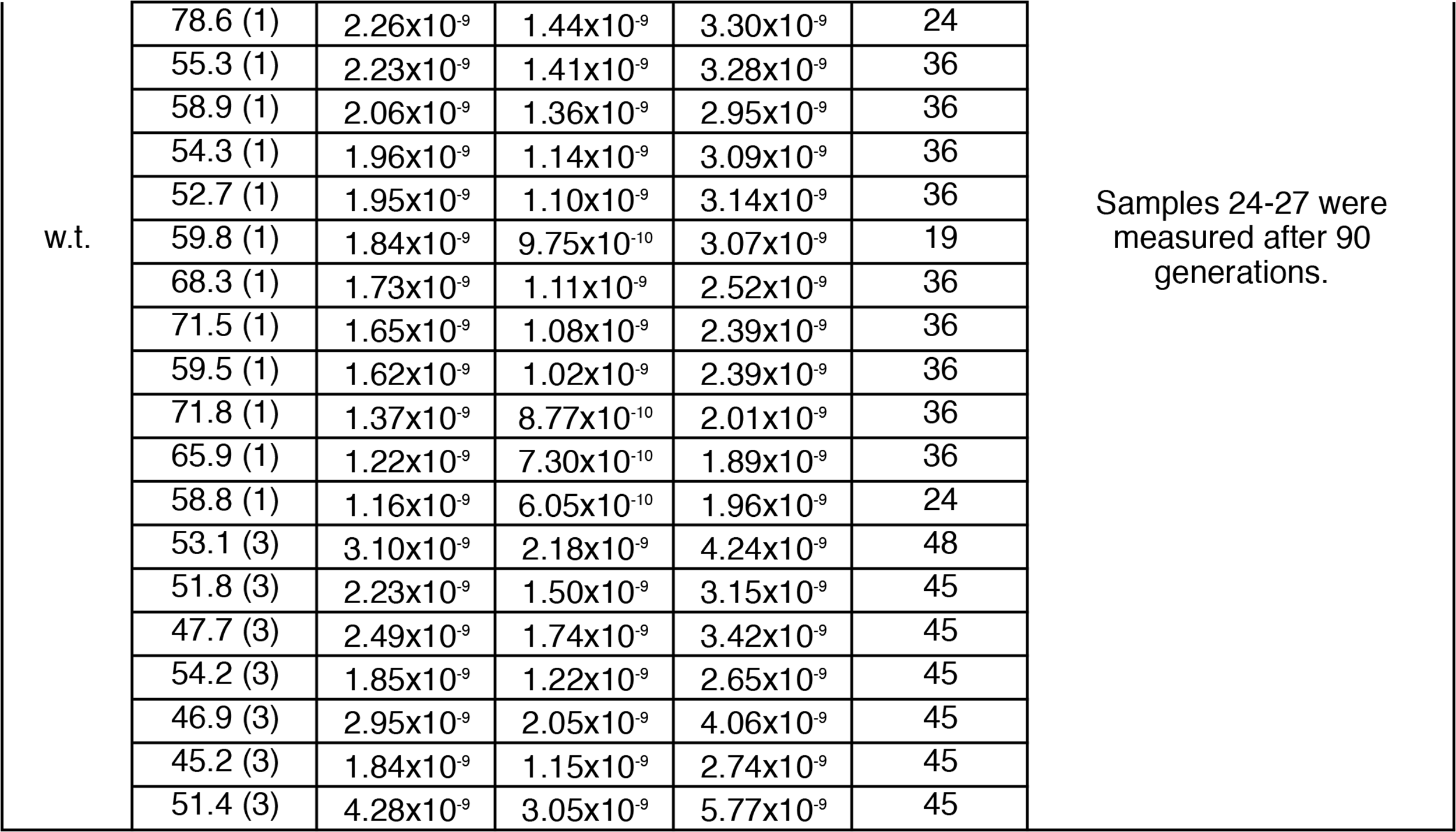
All TP-DNAP1 variants characterized by fluctuation tests in this study. All independent measurements of mutation rate are shown, with corresponding 95% confidence intervals and the number of replicates performed for each fluctuation test listed. The number of replicates assayed for determination of p1 copy number is shown as (n).

**Supplementary Table 3.**
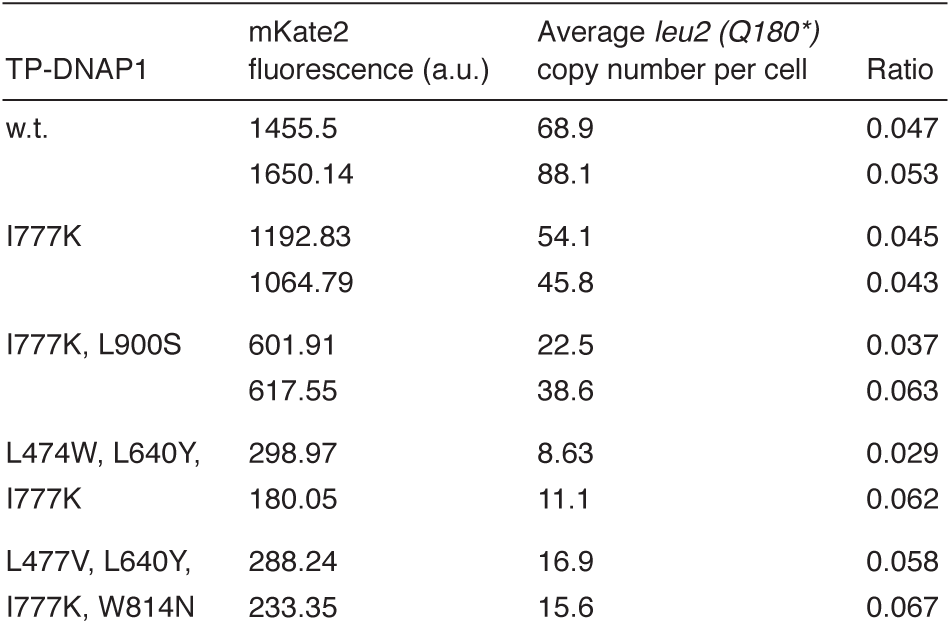
Calibration curve of qPCR-determined p1 copy number to p1-encoded mKate2 fluorescence. Five TP-DNAP1 variants were chosen to represent the range of p1 copy numbers observed across all experiments. OR-Y24 strains containing these TP-DNAP1s were grown in biological duplicates. Each biological duplicate was expanded in technical triplicates for mKate2 fluorescence measurements. In parallel, biological duplicates were expanded in single large volume cultures and subject to DNA extraction and qPCR measurements (see **Methods**). Data shown are mean fluorescence in arbitrary units (a.u.) and qPCR-determined p1 copy numbers. The ratio of copy number to mKate2 fluorescence is shown for each sample. The linear regression has low background and a strong fit (y = 0.048x + 0.206, r^2^ = 0.954).

**Supplementary Table 4.**
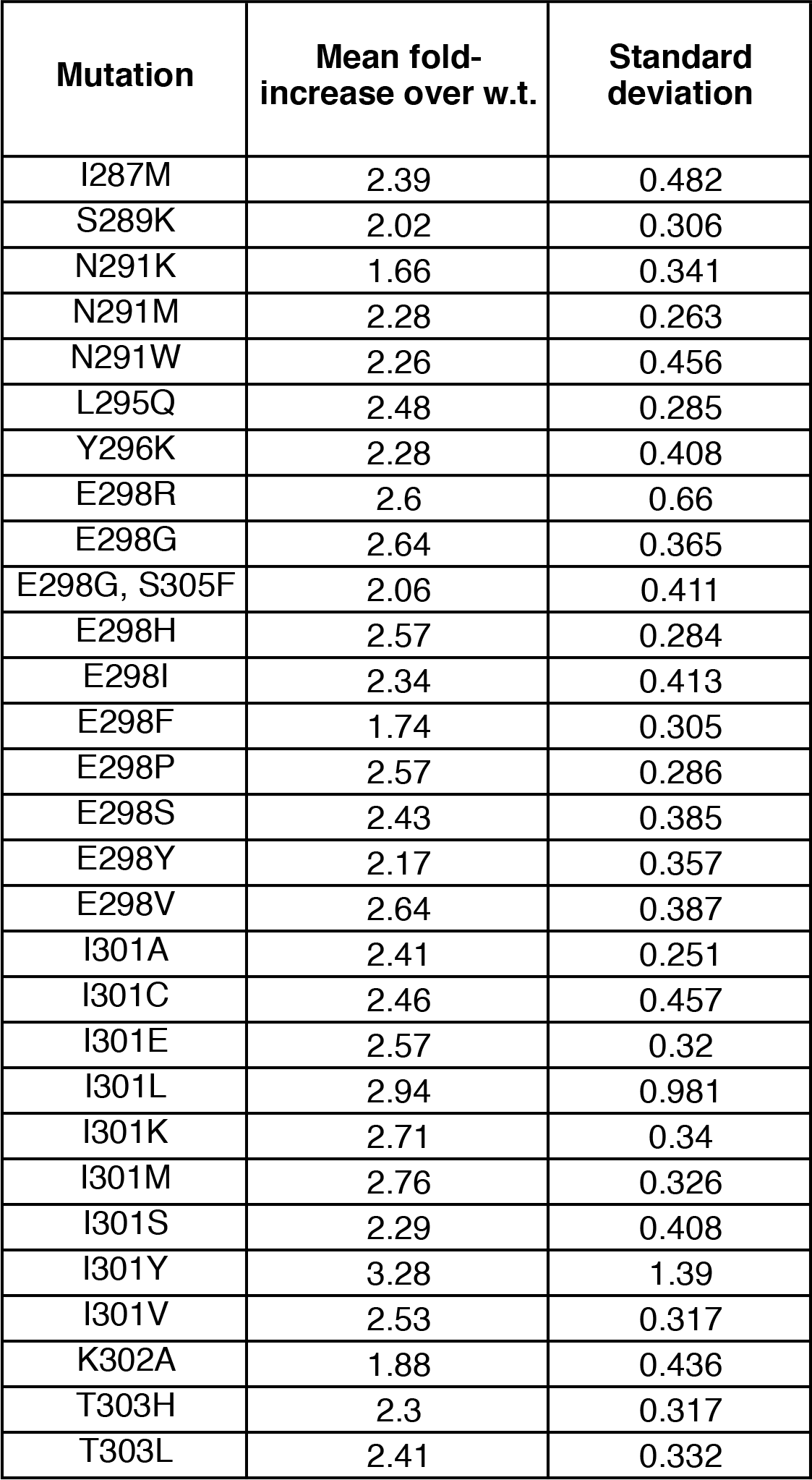
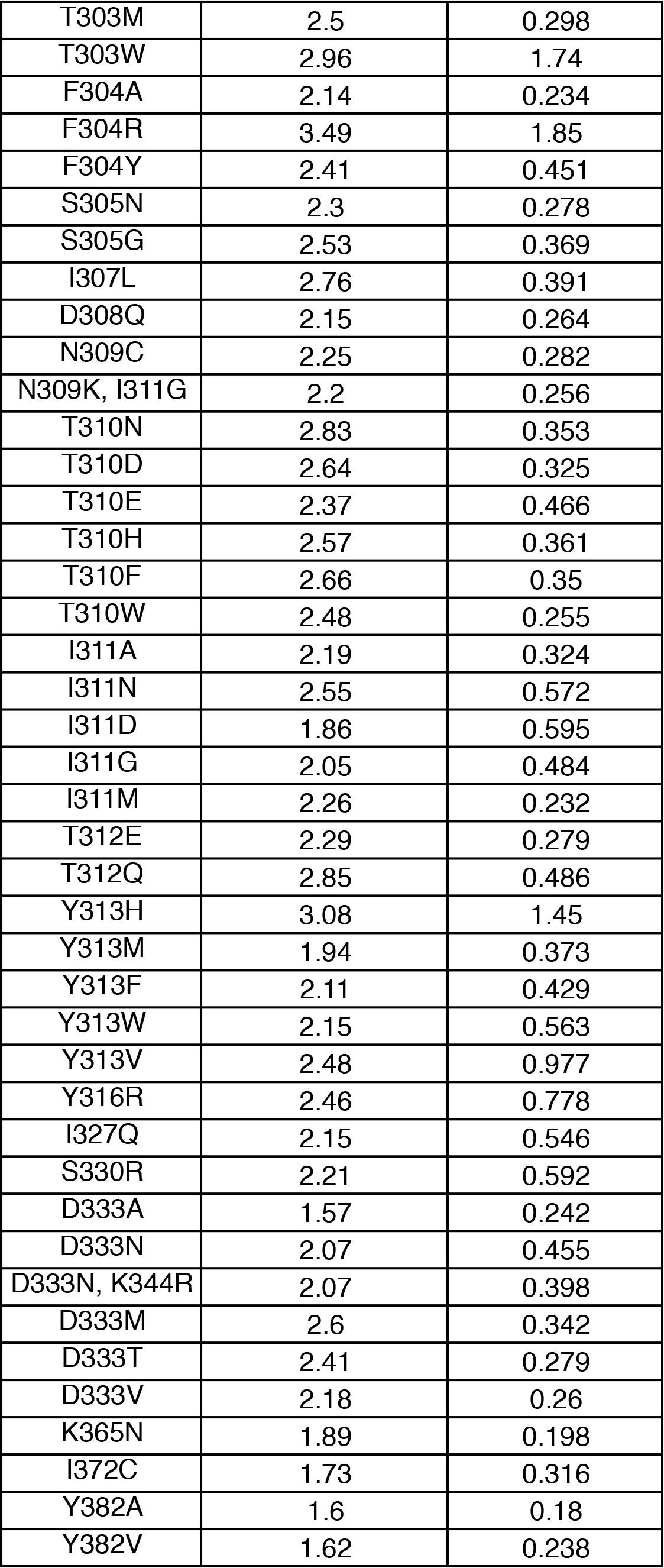
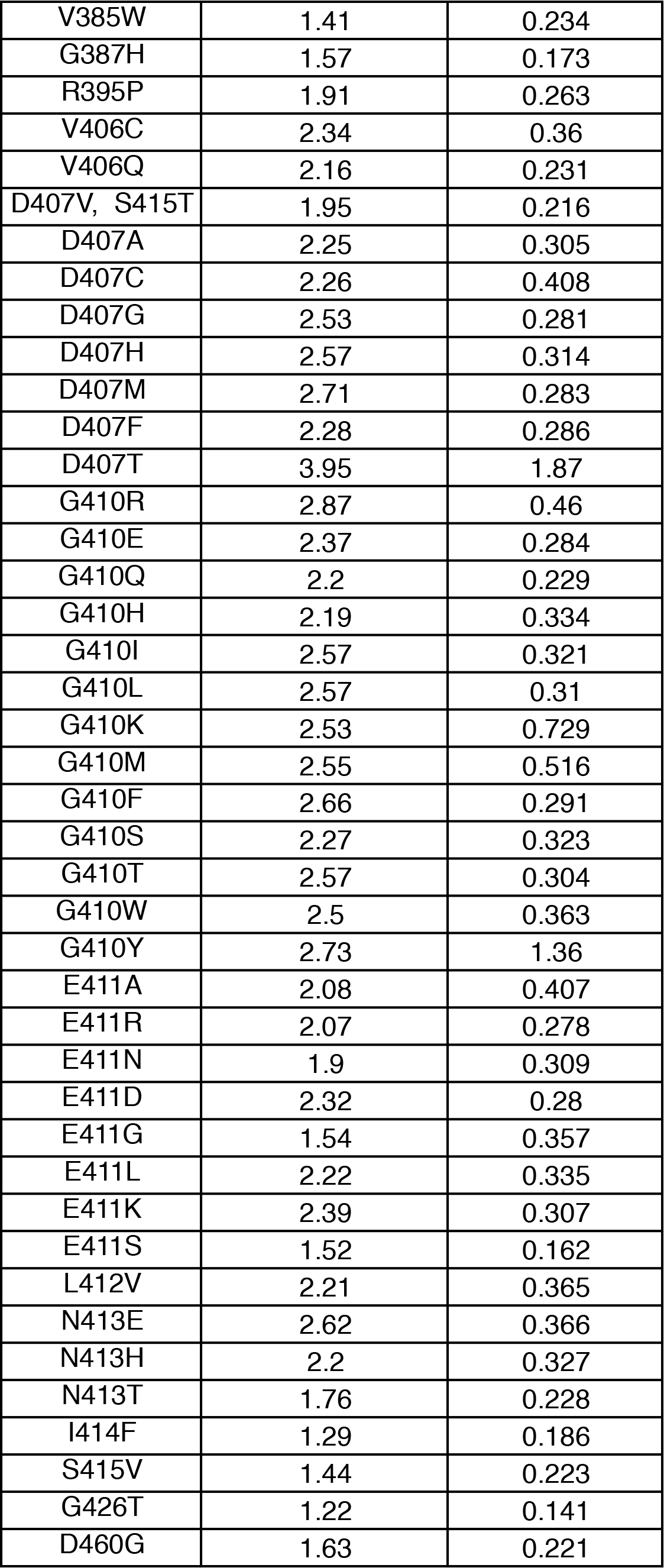
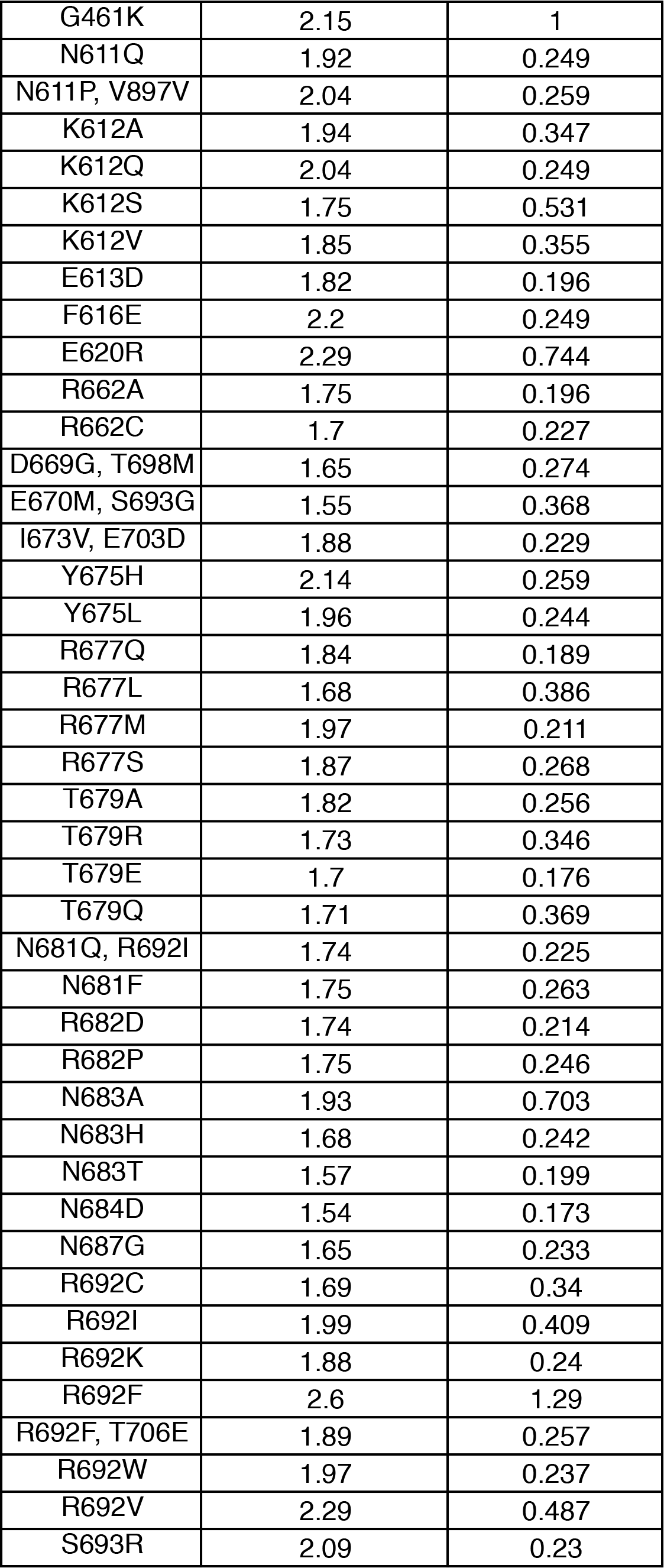
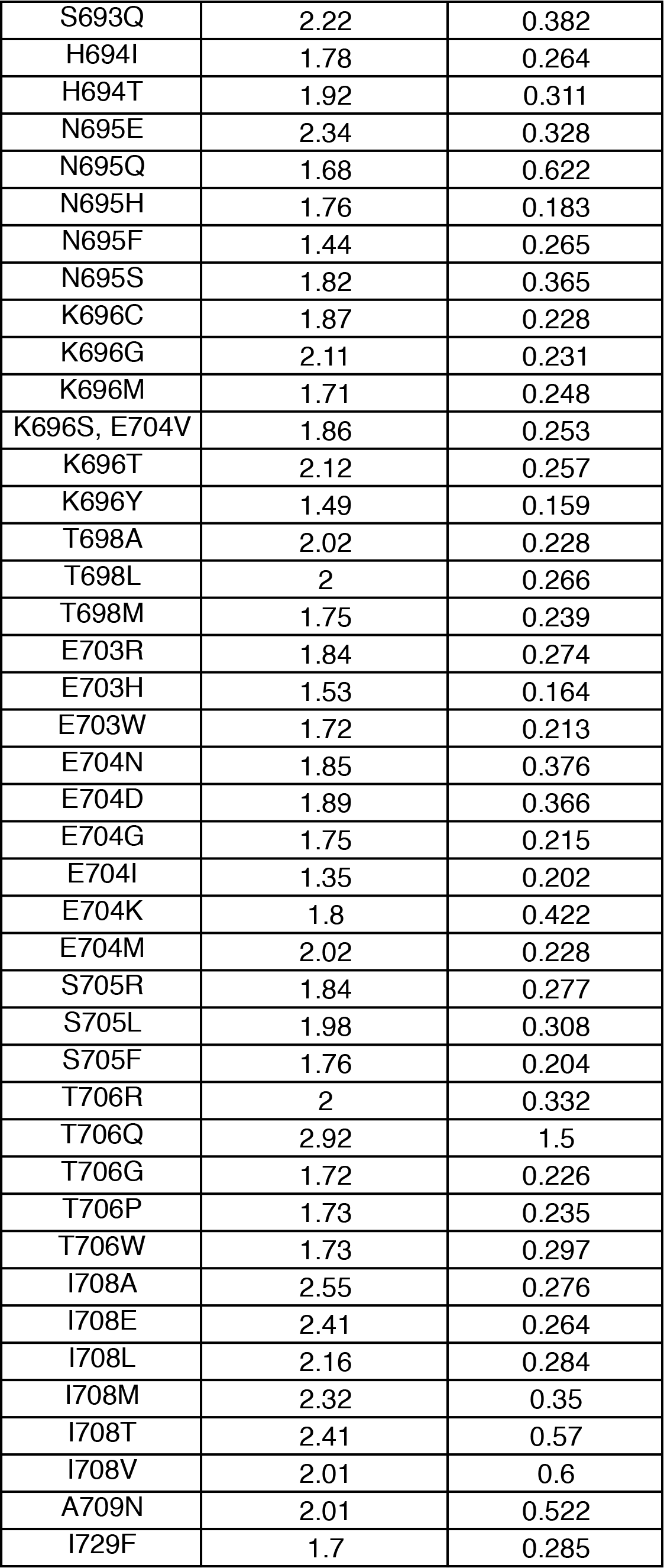
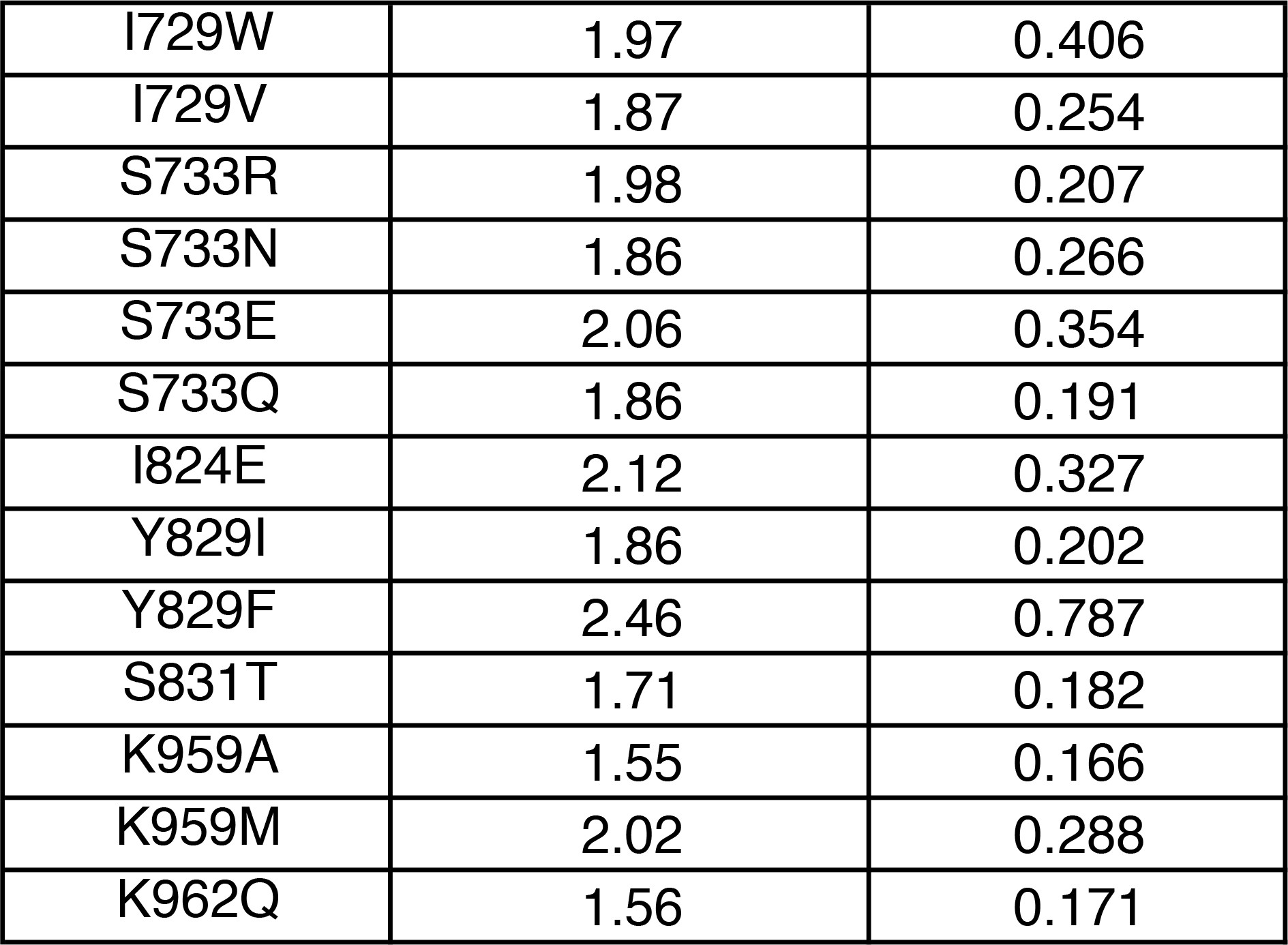
210 TP-DNAP1 variants that replicate p1 at a higher copy number than w.t. TP-DNAP1. From p1 copy number measurements of 13,625 yeast clones screened in small-scale p1 fluctuation tests, 210 unique variants exhibited elevated copy numbers. Variants were re-transformed into OR-Y24 and subject to additional p1 copy number measurements for verification. Data shown are fold-change mean and standard deviation (calculated using equation (5).2 of Frishman^46^) of biological triplicate measurements of each mutant and 15 measurements of w.t. TP-DNAP1. High activity of four TP-DNAP1 variants was independently validated with qPCR measurements of p1 (unpublished results).

**Supplementary Table 5.**
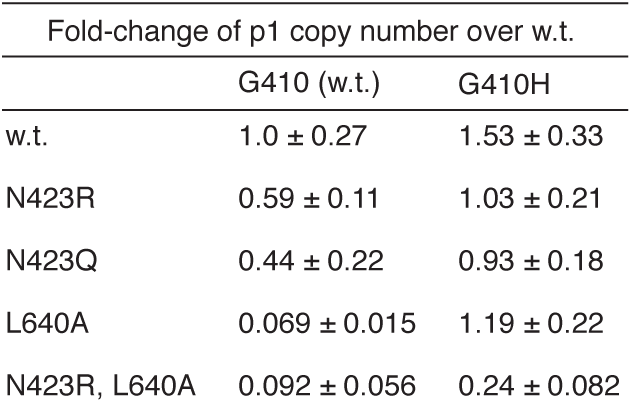
Mutation G410H broadly increases activity of TP-DNAP1 variants. Mutation G410H was added to several low activity TP-DNAP1s. Mutants were transformed into OR-Y24 and subject to p1 copy number measurements. Data shown are mean fold-change ± standard deviation for biological triplicates, calculated using equation (5).2 of Frishman^45^.

**Supplementary Table 6.**
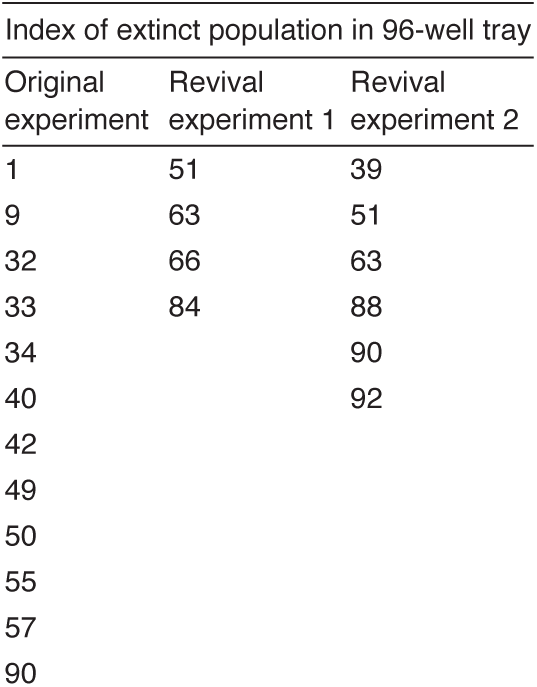
Stochastic extinction in revival experiments of PfDHFR evolution. Indices of populations that went extinct during the 90-replicate *Pf*DHFR evolution experiment and during revival experiments. In the revival experiments, cultures were inoculated from glycerol stocks of passage 5, at which point all 90 populations grew robustly. Cultures were revived in SC media supplemented with 2.5 mM pyrimethamine and passaged with the same protocol used for the original evolution experiment.

**Supplementary Table 7.**
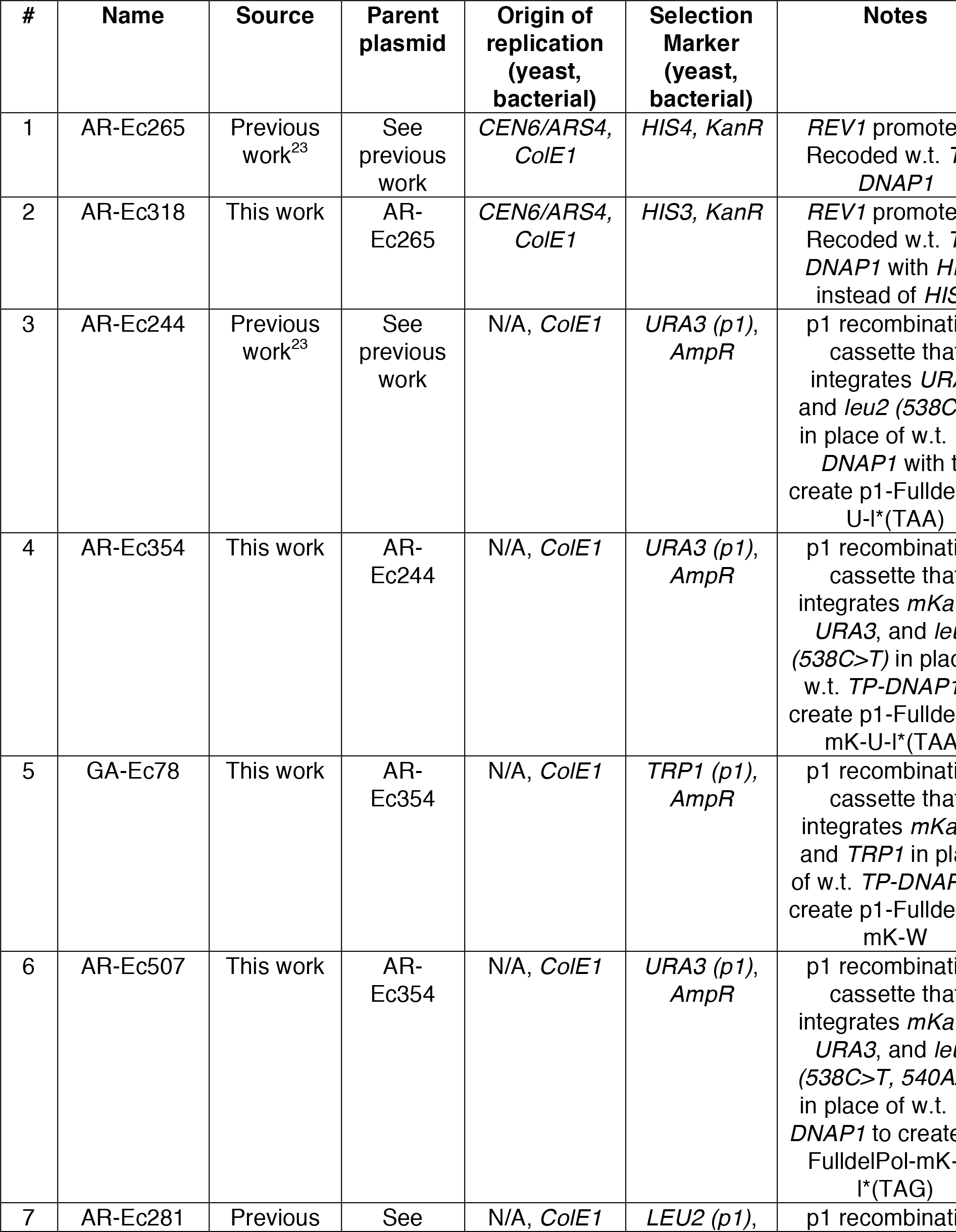
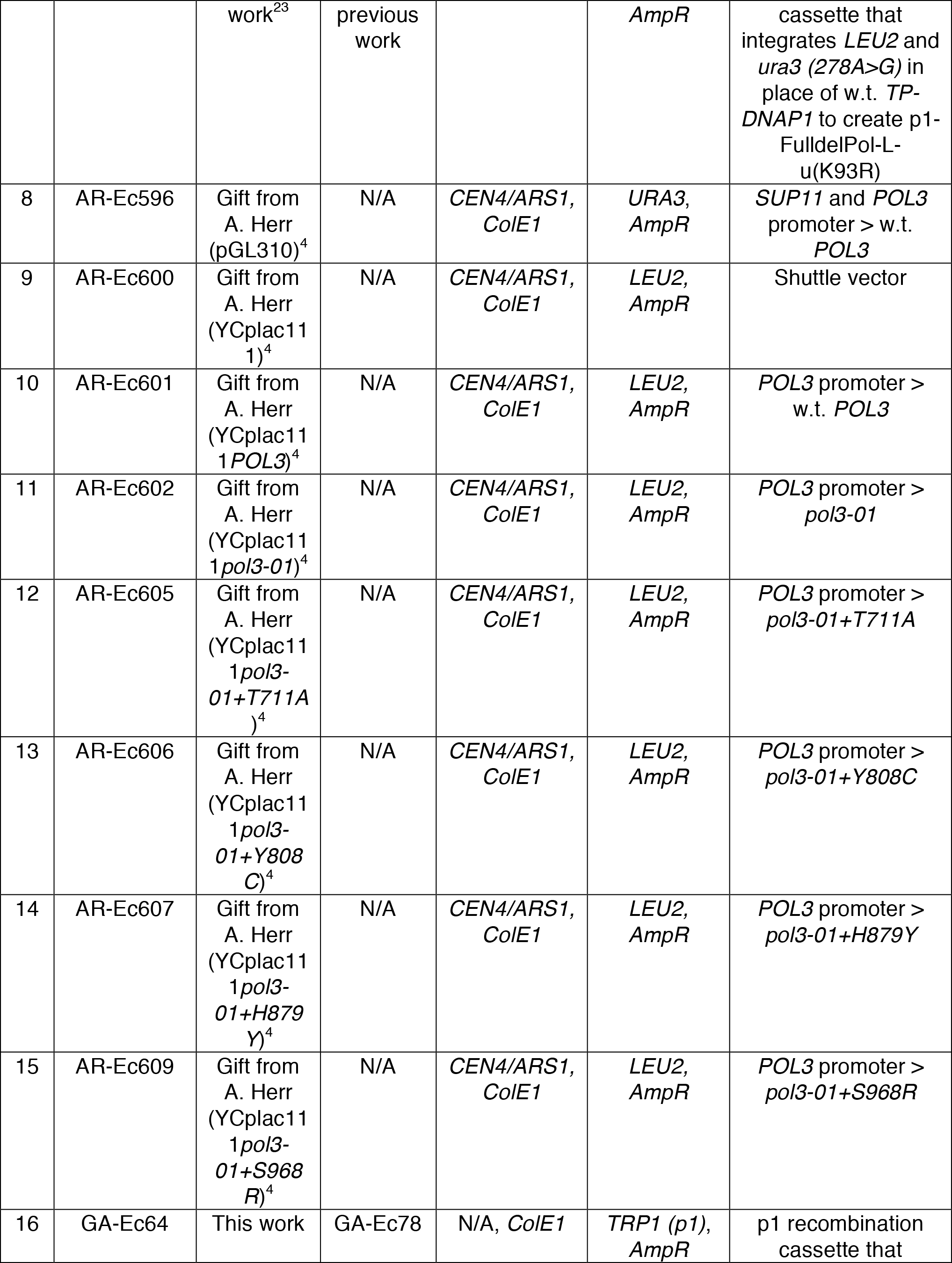
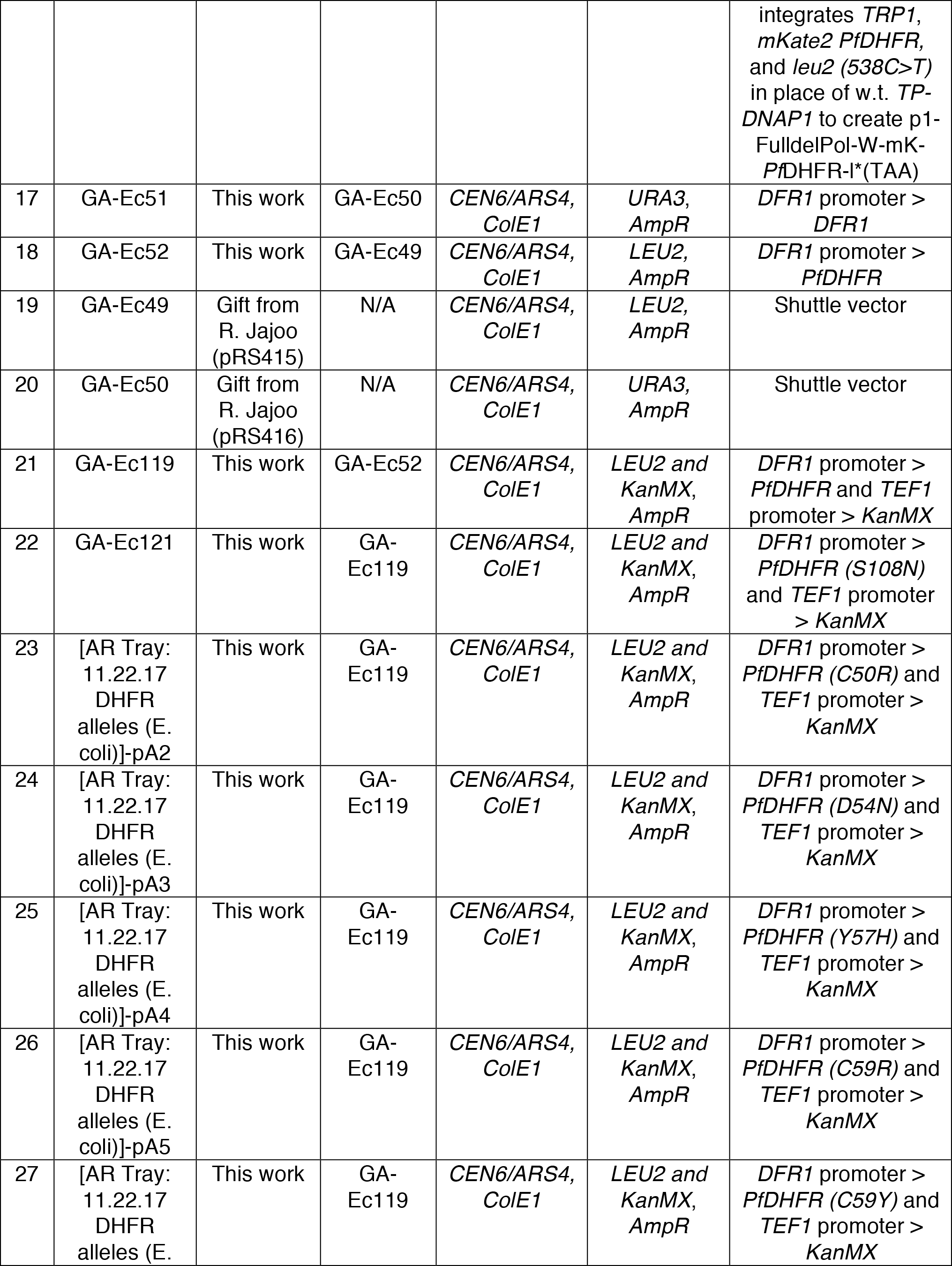
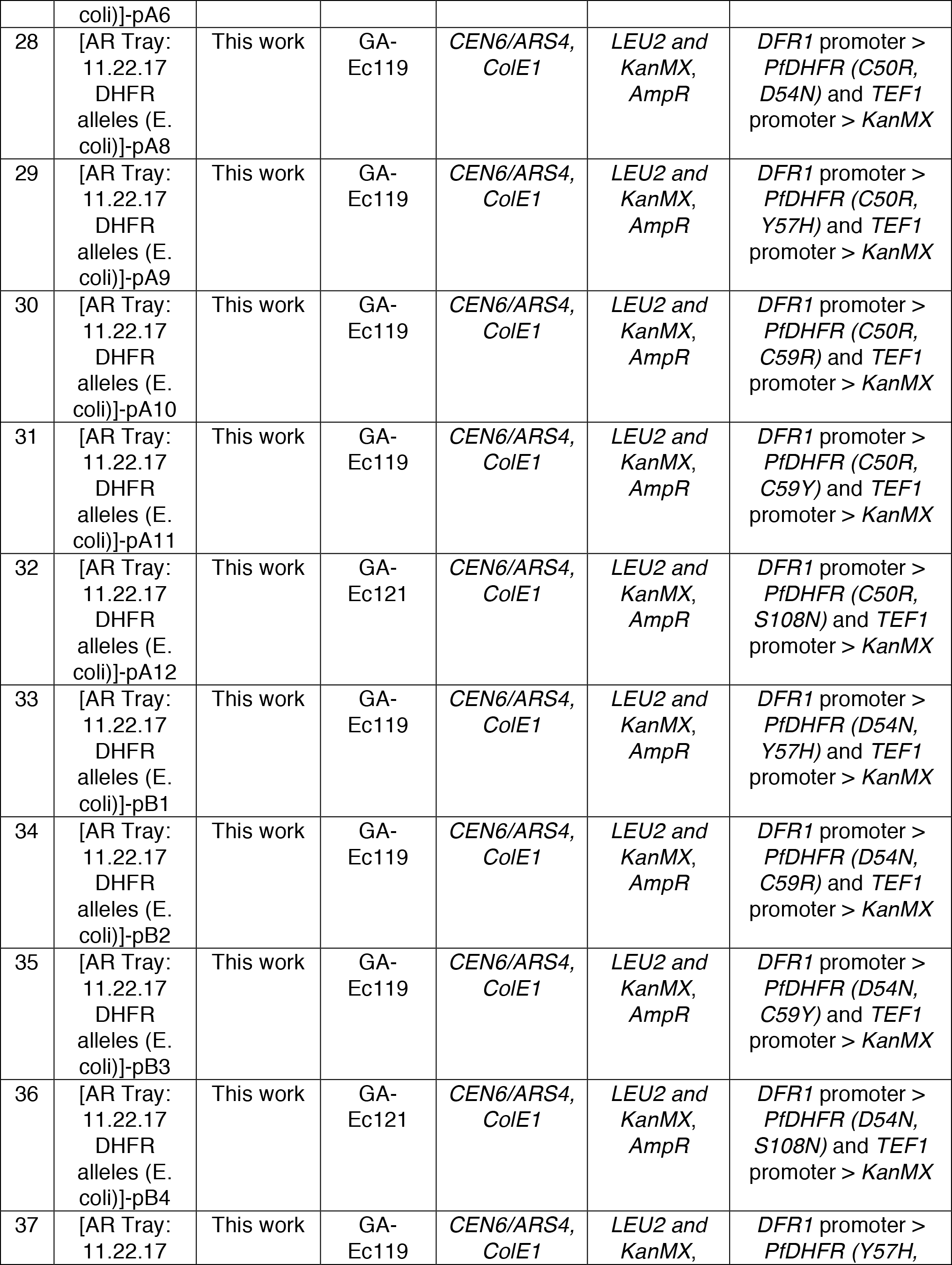
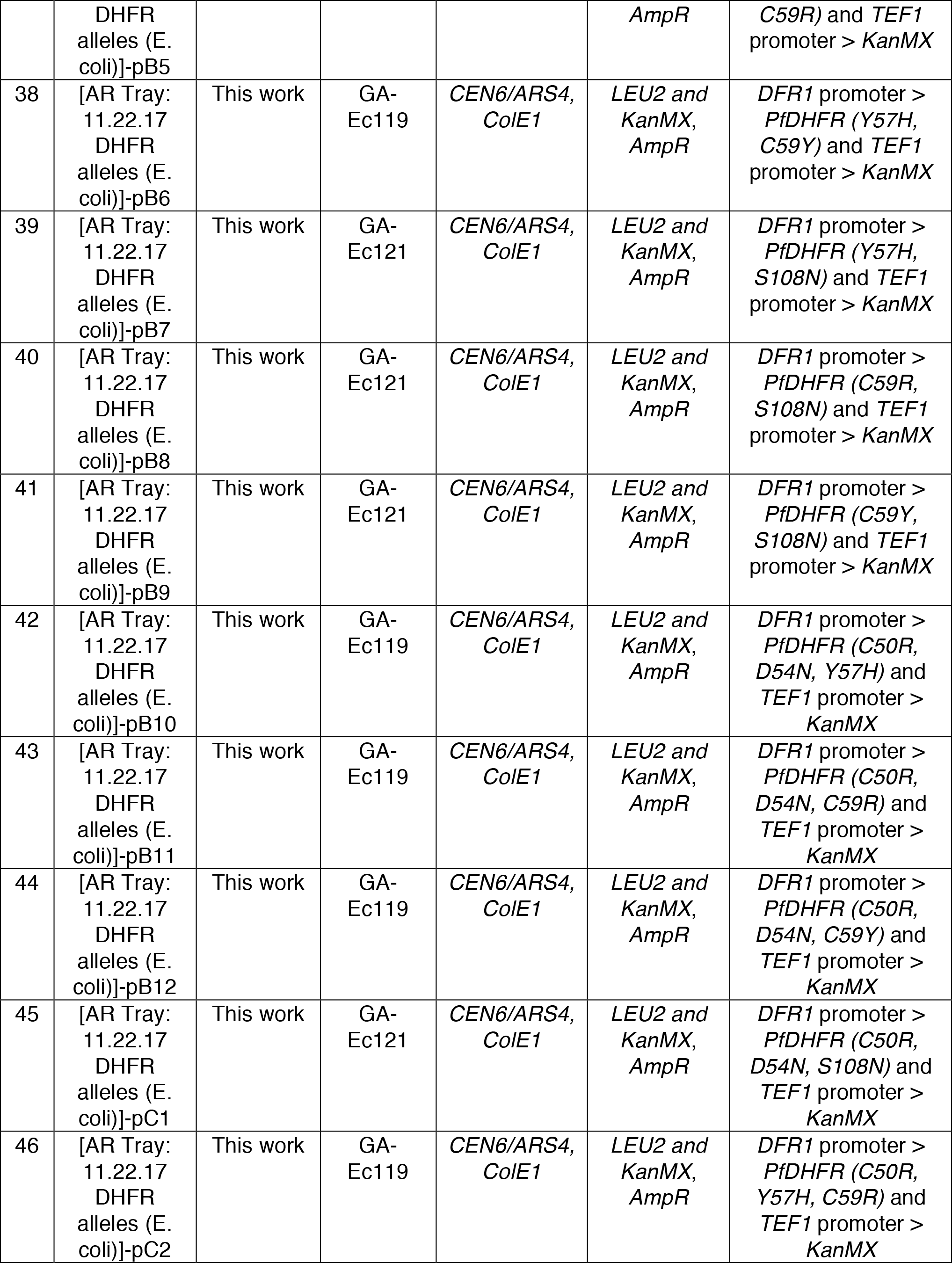
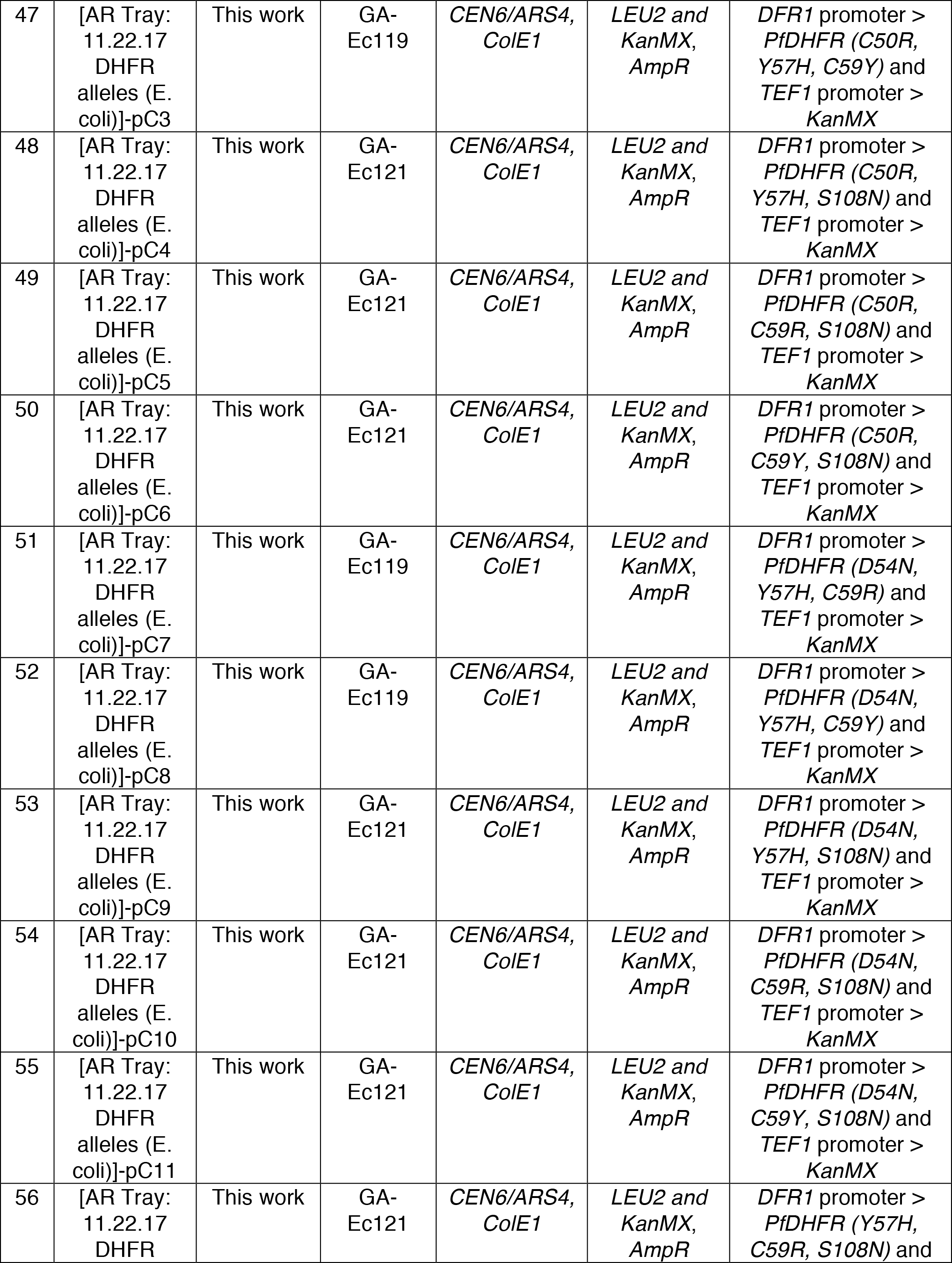
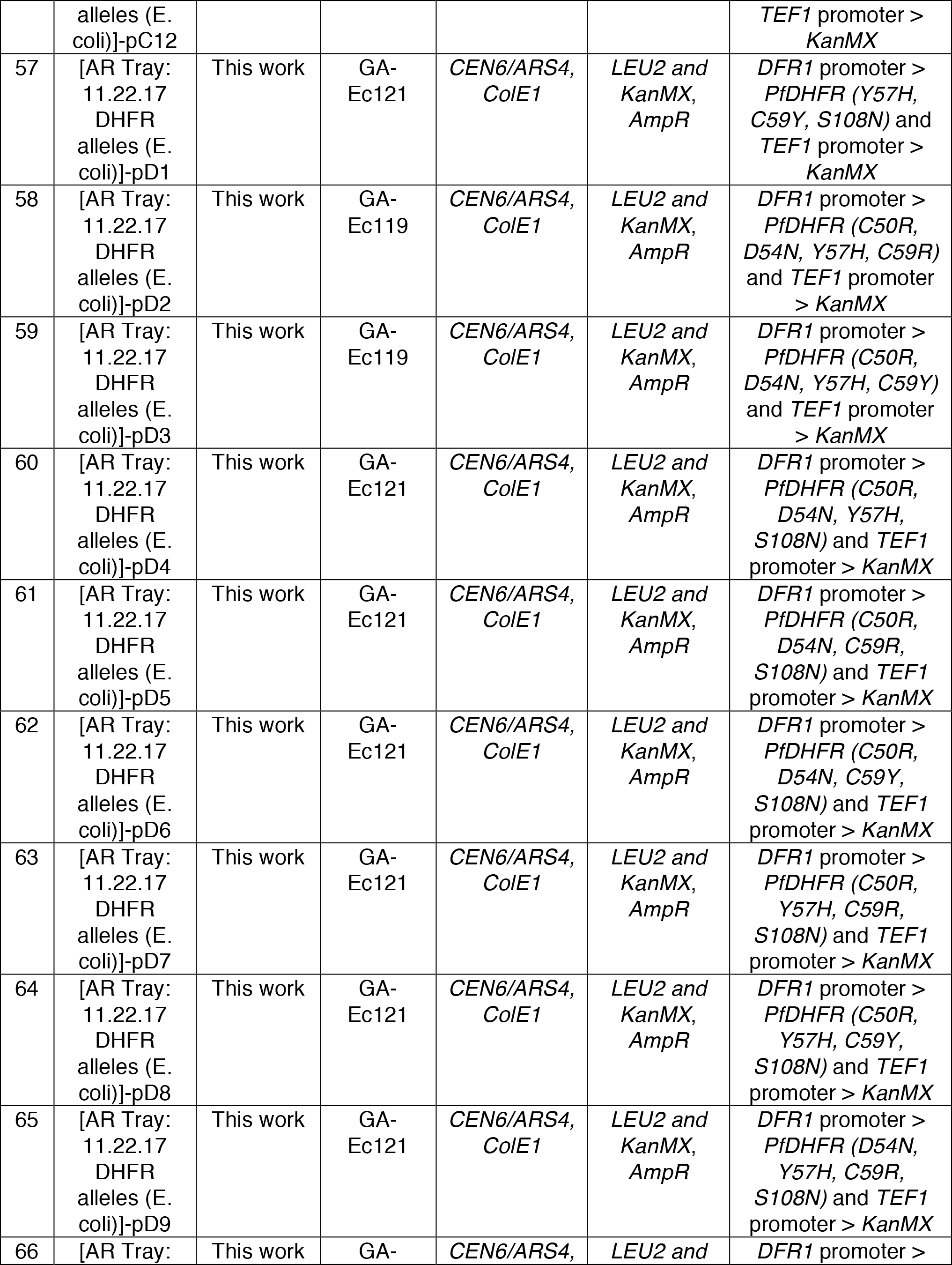
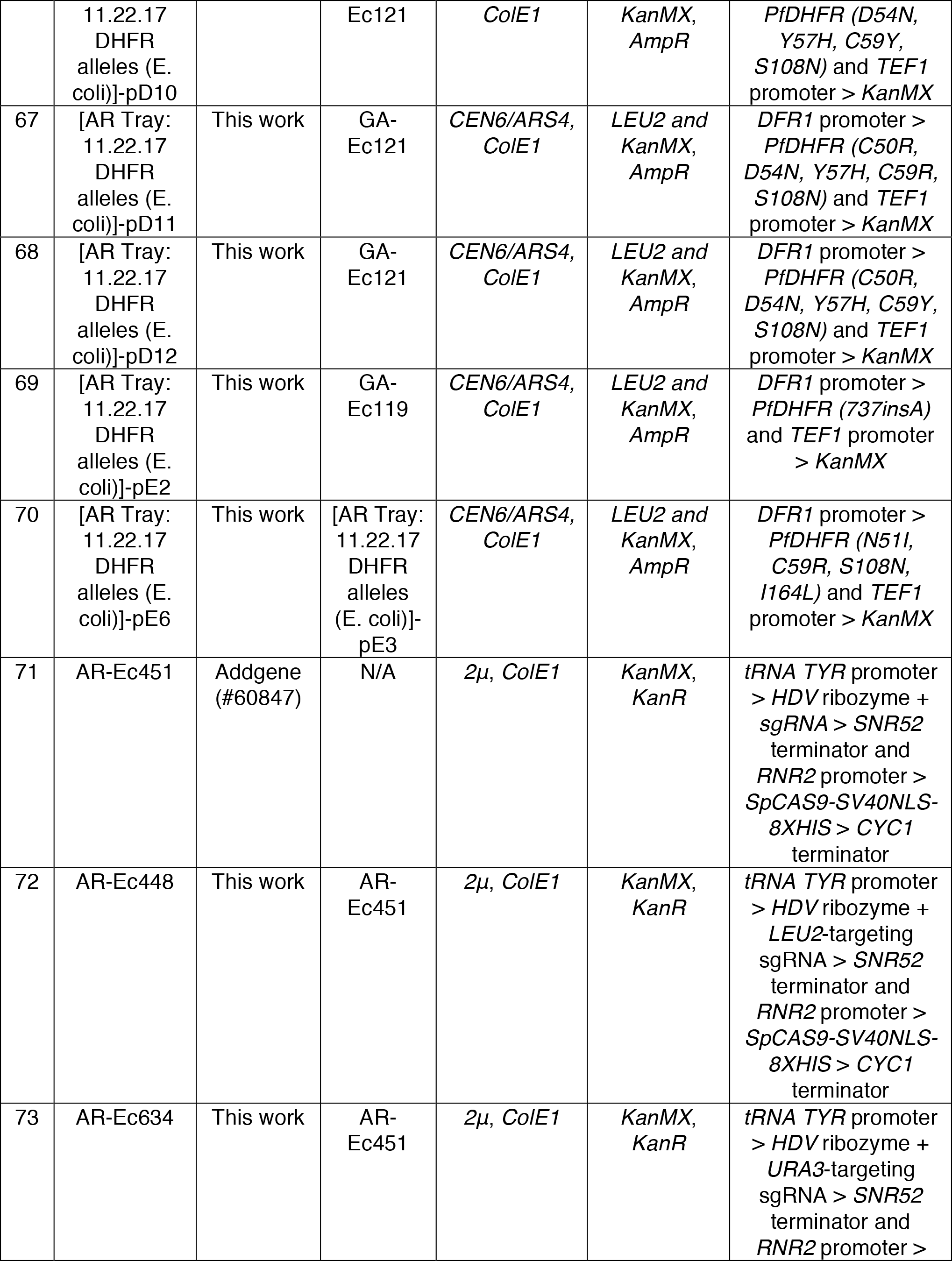
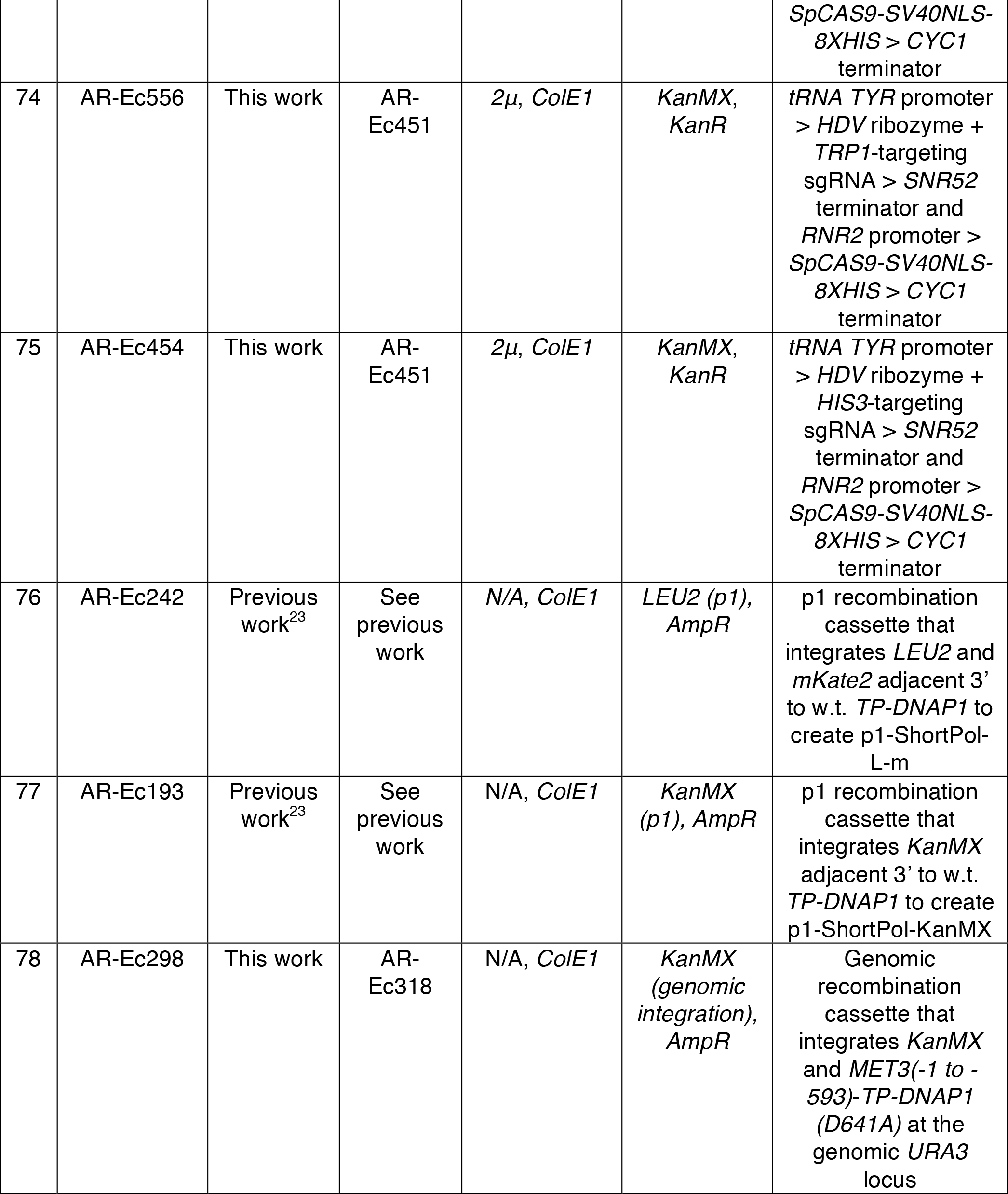
List of plasmid used in this study. Plasmids encoding TP-DNAP1 variants listed in **Supplementary Tables 2, 4**, and **5** are not included. These were all derived from plasmid 2 (described below).

**Supplementary Table 8.**
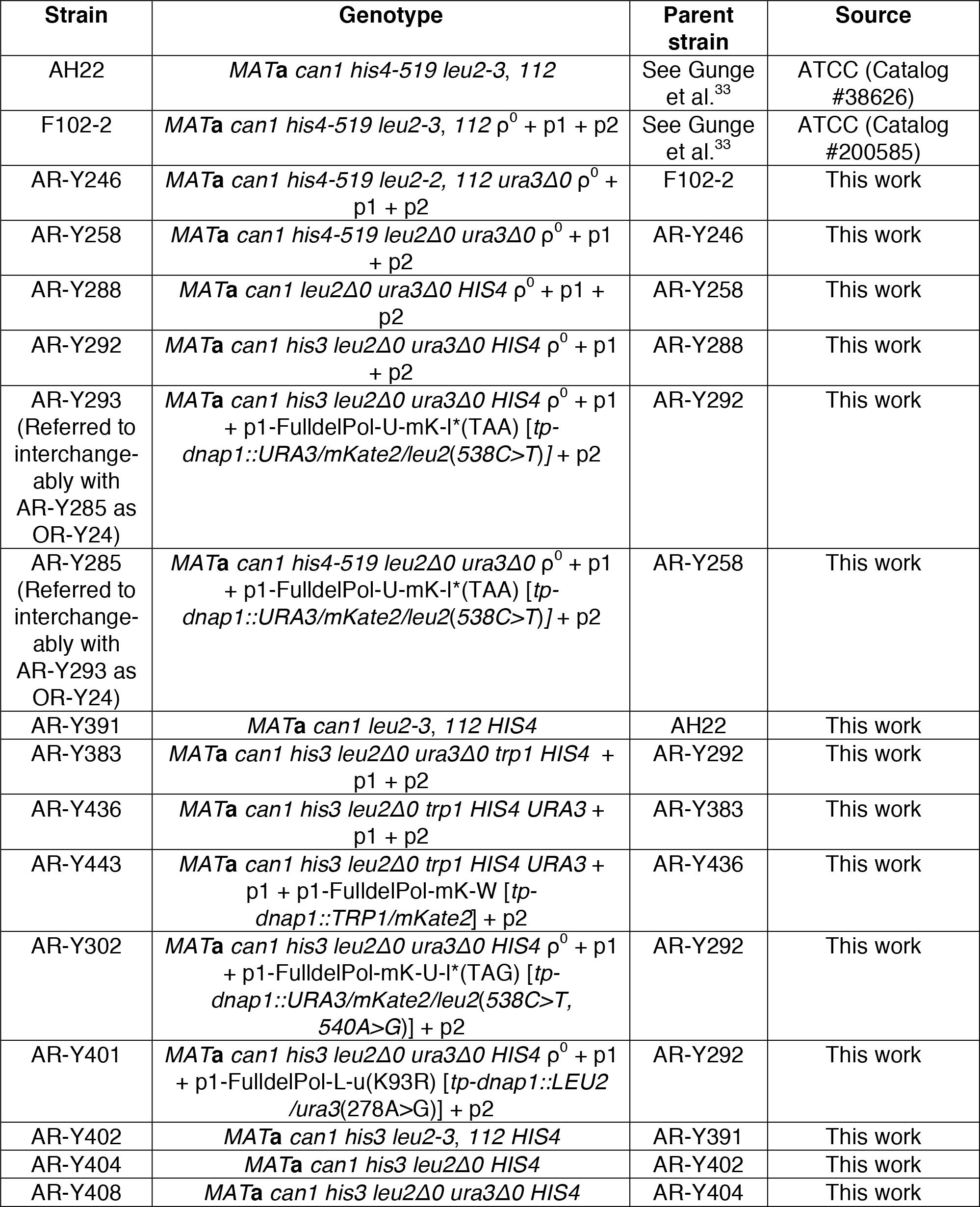
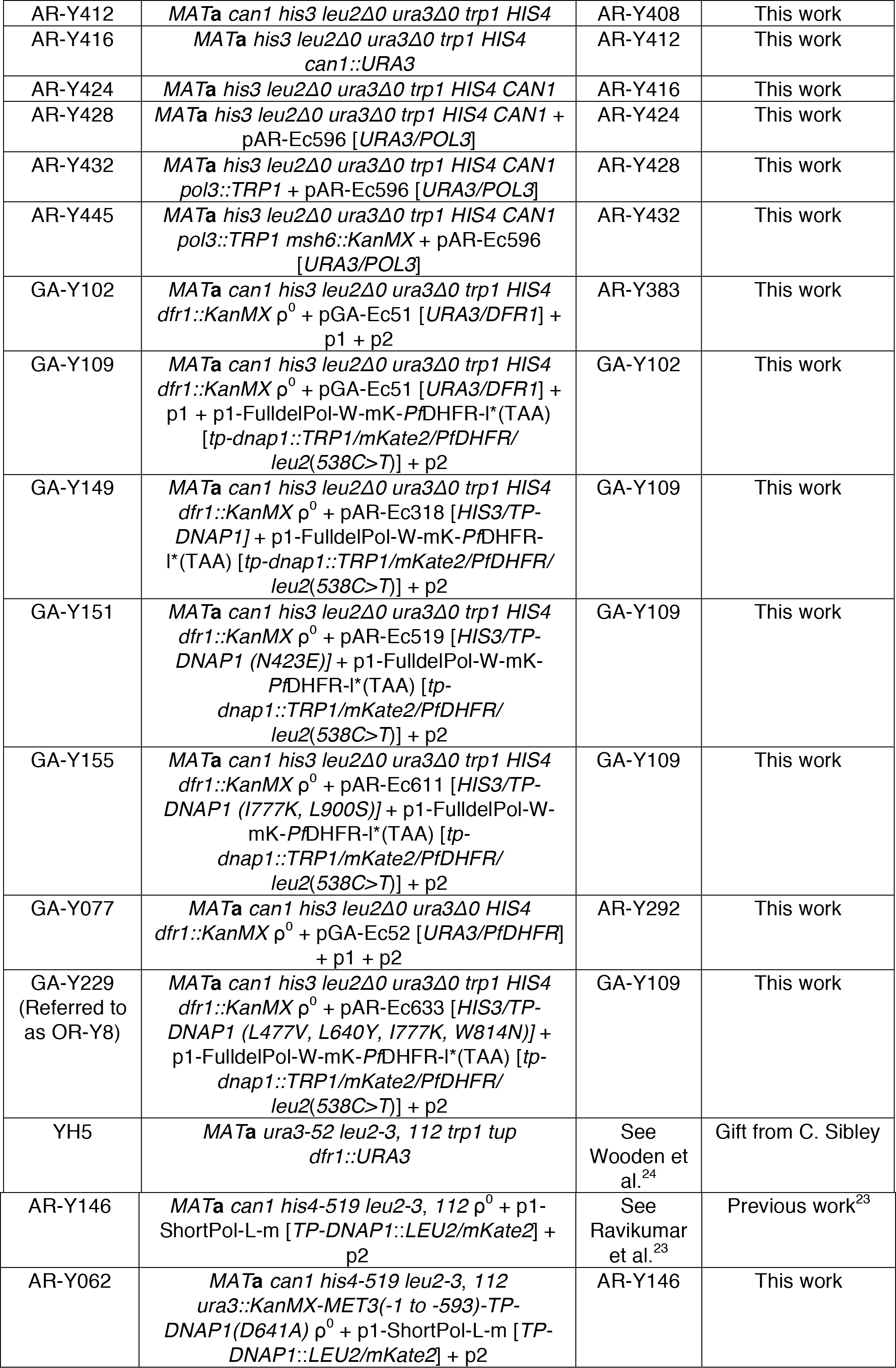
List of parent yeast strains used in this study. Parent strains do not include derivatives of OR-Y24 containing TP-DNAP1 variants, and derivatives of AR-Y432 and AR-Y435 containing POL3 variants.

